# NOMIS: Quantifying morphometric deviations from normality over the lifetime of the adult human brain

**DOI:** 10.1101/2021.01.25.428063

**Authors:** Olivier Potvin, Louis Dieumegarde, Simon Duchesne, the Alzheimer’s Disease Neuroimaging Initiative, the CIMA-Q, the CCNA groups

## Abstract

We present NOMIS (https://github.com/medicslab/NOMIS), a comprehensive open MRI tool to assess morphometric deviation from normality in the adult human brain. Based on MR anatomical images from 6,909 cognitively healthy individuals aged 18-100 years, we modeled 1,344 measures computed using the open access *FreeSurfer* pipeline, considering account personal characteristics (age, sex, intracranial volume) and image quality (resolution, contrast-to-noise ratio and surface reconstruction defect holes), and providing expected values for any new individual. Then, for each measure, the NOMIS tool was built to generate Z-score effect sizes denoting the extent of deviation from the normative sample. Depending on the user need, NOMIS offers four versions of Z-score adjusted on different sets of variables. While all versions consider head size and image quality, they can also incorporate age and/or sex, thereby facilitating multi-site neuromorphometric research across adulthood.

## Introduction

Despite the popularity of magnetic resonance imaging (MRI) to examine abnormalities in brain morphometry, tools quantifying normality are lacking. While age, sex and intracranial volume are well-known to influence brain volume and shape[1, 2] the determination of whether an individual’s brain region measurements are within normality faces multiple major challenges such as the lack of normative data across appropriate age groups, the influence of the MRI processing pipeline, the variety in neuroanatomical atlases used for parcellation and the uniqueness of the image acquisition itself[3, 4]. We made previous attempts[5–8] to produce such normative data in adulthood based on *FreeSurfer*, an open-access and fully automated segmentation software (http://freesurfer.net), for two specific brain atlases, namely Desikan-Killiany[9] (DK) and Desikan-Killiany-Tourville[10] (DKT). This initial foray allowed for the quantification of the extent of deviation from normality for a given individual, according to personal characteristics such as age, sex and estimated intracranial volume (eTIV), while controlling for scanner magnetic field strength (MFS) and original equipment manufacturer (OEM).

Leveraging this prior work, we offer a comprehensive tool called NOMIS (NOrmative Morphometry Image Statistics; https://github.com/medicslab/NOMIS). NOMIS can be used to produce normative values for any new adult individual, cognitively healthy or otherwise. Using this individual’s T1-weighted MRI, processed via the *FreeSurfer* 6.0 toolkit, one can derive Z-score effect sizes denoting the extent of deviation from the normative sample according to the individual’s characteristics (age, sex, and eTIV), while taking into account image quality information (resolution, contrast-to-noise ratio (CNR) and holes in surface reconstruction)[11, 12]. NOMIS contains 1,344 brain measures generated by *FreeSurfer* on 6,909 healthy individuals aged 18 to 100 years (mean ±sd: 55.0 ±20.0; 56.8% female). The normative data includes as before the DK[9] and DKT[10] atlases, as well as the Destrieux (a2009s)[13] neocortical atlas; neocortical pial and white surface areas, volumes and thicknesses; *FreeSurfer*’s default subcortical atlas[14], hippocampal subfields, brainstem subregions; its ex vivo-based labeling protocol atlas[15]; and the subcortical white matter parcellation according to the adjacent neocortical areas. Furthermore, to fulfill specific needs from researchers, we propose four versions of Z-score adjusted on different sets of variables. While all versions are adjusted for head size and image quality, the full version includes both age and sex whereas the three other versions are without age, without sex and without age and sex. Thus, a research group working on aging aiming at removing the variance of hippocampal volumes due to head size, sex, and image quality could use the version without age, which preserves the variance due to aging. When compared to our previous work on normative values, there are important new contributions in NOMIS:

- The norms were calculated using a newer *FreeSurfer* software version
- New variables were added to remove undesirable variance (CNR, surface holes, resolution)
- New atlases were processed, such as Destrieux, hippocampal subfields, brainstem subregions, ex vivo-based labeling protocol atlas, subcortical white matter parcellation according to the adjacent neocortical areas
- The possibility of calculating normative scores while adjusting only some selected variables was introduced (intracranial volume with image quality in combination, or not, with age and/or sex)
- The sample size of the normative sample was doubled, making the age distribution more uniform than previously

### The multiple scanner problem

Different scanners produce different images, even in the same individuals, which produce in the end different morphometric values. One way of capturing inter-scanner variance is using information about the scanner (e.g. magnetic field strength and vendor). For the creation of NOMIS, and contrary to our previous work, we chose not to incorporate such information since the samples of individuals within each combination of scanner characteristic is likely to be different and thus, possibly bias-inducing due to known or unknown individuals’ characteristics stemming from recruitment in a particular study included in the training data. Therefore, to minimize inter-scanner variance, NOMIS strictly uses image information.

Moreover, as a final validation step, we have compared the basic version of NOMIS (i.e. only adjusting for head size and image quality) with two global scaling harmonization techniques, namely NeuroCombat[16] and NeuroHarmonize[17] on their ability to reduce the mean effect and variance induced by different scanners. Such techniques model the differences between scanners to apply a post-hoc correction on morphometric estimates based on the complete set of data in the study. In that, they are fundamentally different from our attempt here at a normative tool to be used in new, separate studies.

Nevertheless, it should be noted that, while they are gaining popularity, harmonization techniques can potentially induce biases due to the different participants’ characteristics at each scanner[18, 19]. The main challenge to show that harmonization is actually working is that MRI provides relative measures for which that there is no gold standard; each scanner yields its own measure, given its hardware software and other factors, even time of the day[20]. In order to properly test harmonization, we defined our own gold standard by using the Single Individual Across Networks (SIMON) dataset[21], comprised of images from a single person that was scanned within a short span at 12 sites for quality control purposes in the context of within two Canadian studies. By harmonizing these 12 scanners using 547 MRIs from individuals scanned in these studies, as well as the quality-control 48 SIMON MRIs, we verified whether the variance of the SIMON measures was lower or not. Unfortunately, we conclude that none of the harmonization techniques reduce real inter-scanner variance. While neither does NOMIS, such is not our purpose.

## Materials and methods

### Normative sample

The norms are based on a cross-sectional sample of 6,909 (initial sample: 7,399) cognitively healthy individuals aged 18 to 100 years, (mean ±sd; 55.0 ±20.0; 56.8% female), gathered from 27 different datasets (Table 1). Supplementary Fig 1 shows the age distribution within each dataset. Scans were acquired from one of the three leading OEM (e.g. Siemens Healthcare (Erlangen, Germany); Philips Medical Systems (Best, Netherlands); or GE Healthcare (Milwaukee, WI)) at MFS of either 1.5 or 3 Tesla. This study received the approval of the Institutional review board of neuroscience and mental health of the CIUSSS de la Capitale-Nationale (#217207).

**Table 1.** Datasets included in the normative sample

| Dataset | n |
| --- | --- |
| Autism Brain Imaging Data Exchange (ABIDE)[23] | 183 |
| Alzheimer's Disease Neuroimaging Initiative (ADNI)[24] | 672 |
| Australian Imaging Biomarkers and Lifestyle flagship study of ageing (AIBL)[25] | 157 |
| Berlin Mind and Brain (Margulies, Villringer) CoRR sample (BMB)[26, 27] | 50 |
| Cambridge Centre for Ageing and Neuroscience (CamCAN)[28, 29] | 630 |
| Center of Biomedical Research Excellence (COBRE)[30] | 70 |
| Cleveland CCF[31] | 30 |
| Consortium for the Early Identification of Alzheimer's Disease (CIMA-Q)[32] | 29 |
| Dallas Lifespan Brain Study (DLBS)[33] | 304 |
| FIND lab sample (FIND) Lab[34] | 13 |
| Functional Biomedical Informatics Research Network (FBIRN)[35] | 33 |
| Lifespan Human Connectome Project in Aging (HCP-Aging)[36] | 612 |
| International Consortium for Brain Mapping (ICBM) - MNI[37] | 147 |
| Information eXtraction from Images (IXI)[38] | 554 |
| F.M. Kirby Research Center neuroimaging reproducibility data (KIRBY-21)[39] | 20 |
| Minimal Interval Resonance Imaging in Alzheimer's Disease (MIRIAD)[40] | 21 |
| National Alzheimer's Coordinating Center (NACC)[41] | 1562 |
| National Database for Autism Research (NDAR)[42] | 56 |
| Nathan Kline Institute Rockland Sample(NKI-RS)[43] | 138 |
| Nathan Kline Institute Rockland Enhanced Sample (NKI-RES) [43] | 436 |
| Open Access Series of Imaging Studies (OASIS)[44] | 288 |
| POWER Neuroimage sample (POWER)[45] | 26 |
| Parkinson's Progression Markers Initiative (PPMI)[46] | 158 |
| Southwest University Adult Lifespan Dataset (SALD)[47] | 490 |
| University of Wisconsin (Birn, Prabhakaran, Meyerand) CoRR sample (UWM)[26] | 25 |
| Wayne State EF Dataset (Wayne State)[48] | 108 |
| Yale Low-Resolution Controls Dataset (Yale)[49] | 97 |
| Total | 6909 |

From all the samples mentioned, only cognitively healthy (control) participants were included. For the Nathan Kline Institute samples, which were projects recruiting in the general population, we excluded participants with history of schizophrenia or other psychotic disorders, bipolar disorders, major depressive disorders (recurrent), posttraumatic stress disorder, substance abuse/dependence disorders, neurodegenerative and neurological disorders, head injury with loss of consciousness/amnesia, and lead poisoning. Moreover, for the Parkinson’s Progression Markers Initiative dataset, we excluded participants with a Geriatric Depression Scale[22] score of more than 5.

Among the datasets are the Alzheimer’s Disease Neuroimaging Initiative (ADNI), the Australian Imaging, Biomarkers and Lifestyle study of aging (AIBL) and the Consortium pour l’identification précoce de la maladie Alzheimer - Québec (CIMA-Q) datasets. The ADNI (adni.loni.usc.edu) was launched in 2003 as a public-private partnership, led by Principal Investigator Michael W. Weiner, MD. (www.adni-info.org). The AIBL data was collected by the AIBL study group and AIBL study methodology has been reported previously by Ellis et al. (2009). For each dataset, approval from the local ethics board and informed consent of the participants were obtained. Founded in 2013, the main objective of CIMA-Q is to build a cohort of participants characterized in terms of cognition, neuroimaging and clinical outcomes in order to acquire biological samples allowing (1) to establish early diagnoses of Alzheimer’s disease, (2) to provide a well characterized cohort and (3) to identify new therapeutic targets. The principal investigator and director of CIMA-Q is Dr Sylvie Belleville from the Centre de recherche de l’Institut universitaire de gériatrie de Montréal, CIUSSS Centre-sud-de-l’Île-de-Montréal. CIMA-Q represent a common effort of several researchers from Québec affiliated to Université Laval, Université McGill, Université de Montréal, et Université de Sherbrooke. CIMA-Q recruited cognitively healthy participants, participants with subjective cognitive impairment, mild cognitive impairment, or Alzheimer’s disease, between 2013–2016. Volunteers were recruited from memory clinics, through advertisements posted in the community and amongst participants from the NuAge population study.

### Harmonization test sample

For the harmonization test, we used three datasets: 1) the complete CIMA-Q sample (n=208 participants; 286 MRIs), which was described earlier in the method section, 2) the Comprehensive Assessment of Neurodegeneration and Dementia (COMPASS-ND; n=393) study conducted by the Canadian Consortium on Neurodegeneration in Aging (CCNA), and the 3) the SIMON dataset[19], comprised of images from single healthy volunteer scanned on the same scanner as those used for CIMA-Q and COMPASS-ND. From COMPASS-ND, we used participants that were either cognitively unimpaired participants (CU), with mild cognitive impairment (MCI), and with probable Alzheimer’s disease (AD), for a total of 273 participants. While CIMA-Q and COMPASS-ND were acquired at 18 different sites, we selected only data from scanners that had at least three participants other than SIMON, which resulted in a total of 547 images (300 CU, 193 MCI, 54 AD) from 12 different scanners; each ranging from 7 to 145 participants. On those 12 scanners, SIMON was scanned 48 times and was aged between 42-46 years old during that time.

### Brain segmentation

Brain segmentation was conducted using *FreeSurfer* version 6.0, a widely used and freely available automated processing pipeline that quantifies brain anatomy (http://freesurfer.net). All raw T1-weighted images were processed using the “recon-all-all” *FreeSurfer* command with the fully-automated directive parameters (no manual intervention or expert flag options) on the CBRAIN platform[50]. Normative data were computed for volumes, neocortical thicknesses and white and pial surfaces areas for all atlases comprised in *FreeSurfer* 6.0: the default subcortical atlas[14] (aseg.stats), the Desikan-Killiany atlas[9] (DK, aparc.stats file), the Desikan-Killiany-Tourville atlas[10] (DKT, aparc.DKT.stats file), the Destrieux atlas[13] (aparc.a2009s.stats file), the ex vivo atlas,[51] including entorhinal and perirhinal cortices, the brainstem sub-regions atlas[52], the Brodmann area maps which includes somatosensory areas, several motor and visual areas, as well as the hippocampal subfields atlas[53].

The technical details of *FreeSurfer*’s procedures are described in prior publications. Briefly, this processing includes motion correction, removal of non-brain tissue using a hybrid watershed/surface deformation procedure, automated Talairach transformation, intensity normalization, tessellation of the gray matter white matter boundary, automated topology correction, and surface deformation following intensity gradients to optimally place the gray/white and gray/cerebrospinal fluid borders at the location where the greatest shift in intensity defines the transition to the other tissue class. Once the cortical models are complete, a number of deformable procedures can be performed for further data processing and analysis including surface inflation, registration to a spherical atlas which is based on individual cortical folding patterns to match cortical geometry across subjects and parcellation of the cerebral cortex into units with respect to gyral and sulcal structure. This method uses both intensity and continuity information from the entire three-dimensional MRI volume in segmentation and deformation procedures to produce representations of cortical thickness, calculated as the closest distance from the gray/white boundary to the gray/cerebrospinal fluid boundary at each vertex on the tessellated surface. The maps are created using spatial intensity gradients across tissue classes and are therefore not simply reliant on absolute signal intensity. Procedures for the measurement of cortical thickness have been validated against histological analysis [54] and manual measurements[55, 56]. Estimated intracranial volume[57] was taken from the aseg.stats *FreeSurfer* output file. We added the total ventricle volume (labeled as “ventricles”) using the sum of all ventricles and the corpus callosum (labeled as “cc”) using the sum of all corpus callosum segments.

### Quality control and sample selection

A flow chart detailing the final analysis sample is shown in Fig 1. From an initial pool of 7,399 MRIs, nine images failed the *FreeSurfer* pipeline. Following processing, each of the remaining 7,390 brain segmentations was visually inspected through at least 20 evenly distributed coronal sections by O.P. (see Supplementary materials for quality control examples). After quality control, 445 images (6.0%) were removed from further analyses due to segmentation problems, the main reason being that parts of the brain were not completely segmented (e.g. temporal and occipital poles. During visual inspection, 26 brains were found to have signal alterations or clear significant brain lesions and were excluded. Quality control image examples are displayed as Supplementary materials (setup, segmentation errors and abnormalities). In addition to visual inspection, we excluded participants if at least one of the 1,344 brain region measures was missing (n=10). In fine, the analysis sample was composed of 6,909 individual MRIs.

**Fig 1.**
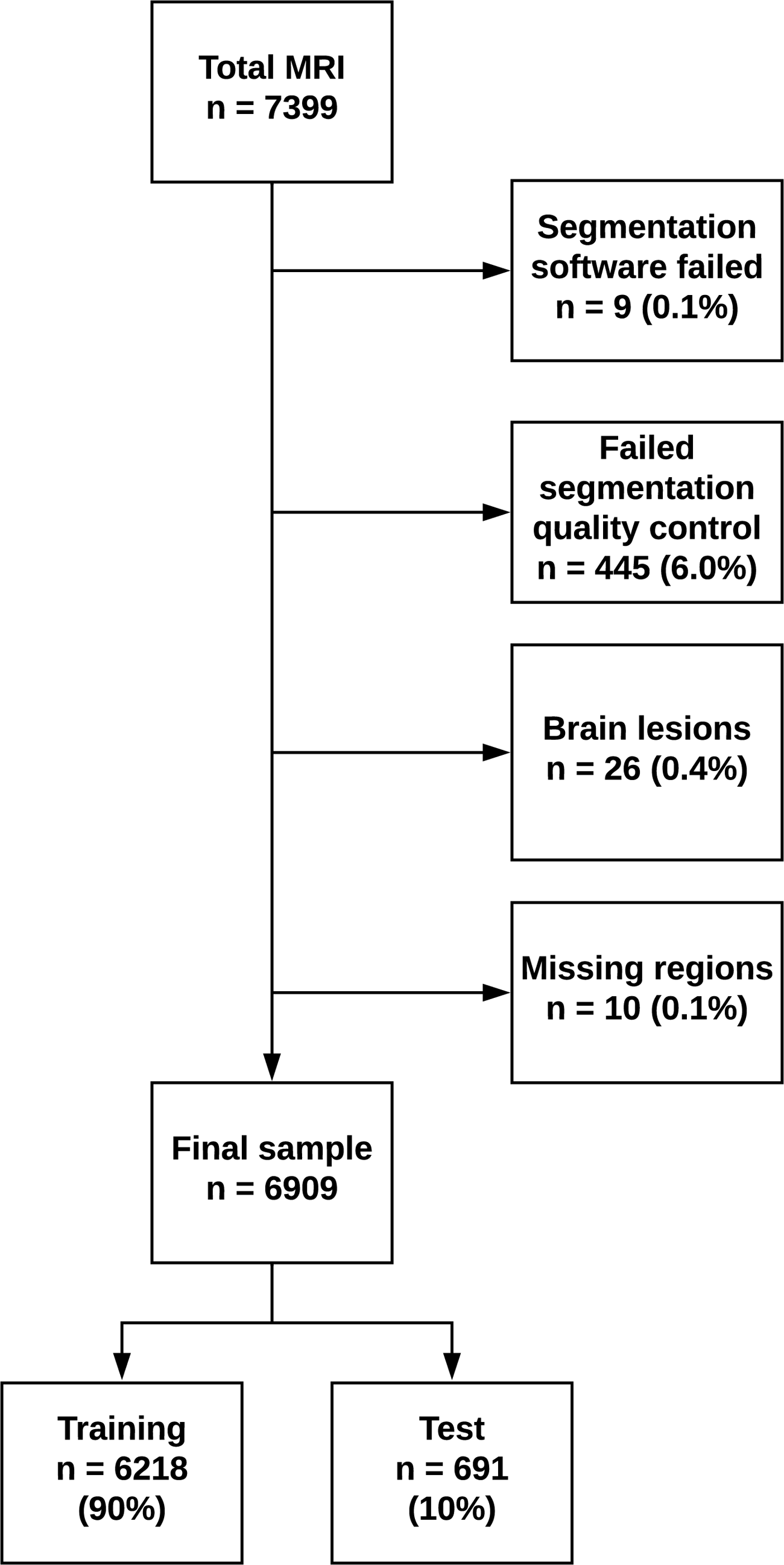
Flow chart of the images.

### Training, validation and test sample

We randomly selected 10% of the whole sample (n=691) to test the models in an independent sample (age: 55.1 ±20.1, range 18-100; 58.5% female). This test sample was not used to build the models predicting normative values. The remaining 90% was used as training sample (age: 54.9 ±20.0, range 18-100; 56.7% female) to build and validate the models. Leave-10%-out cross-validation was used to validate the model in the training sample.

### Clinical samples evaluations

We evaluated the usefulness of normative values using clinical samples of individuals with schizophrenia (n=72; Age: 38.2 ±13.9, range 18-65; 19% female) from the COBRE dataset, as well as participants with clinically ascertained Alzheimer’s disease (n=157 Age: 74.8 ±8.1, range 55-90; 43% female) from the baseline ADNI-2 dataset.

### Image quality predictors

Image quality predictors included voxel size (resolution) and two measures of image quality, one global, and the second local. The first was the total number of defect holes over the whole cortex, i.e. topological errors in the initial cortical surface reconstructions. The total number correlated well with visual inspection of the whole image by trained raters [11]. This measure was extracted from the aseg.stats *FreeSurfer* output file. The second measure was contrast-to-noise ratio (CNR) assessed in each region (R) and therefore used as a regional measure of image quality. For each region, CNR was calculated after *FreeSurfer* preprocessing using gray matter (GM) and cerebral white matter (WM) intensities from the brain.mgz file and the following formula:

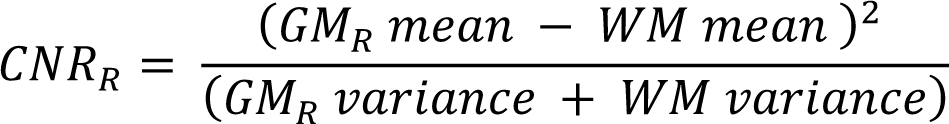

### Outliers removal

For each brain measure, to exclude potential abnormalities, outliers with Z scores lower than −3.29 and higher than 3.29 (*p* < .001) were removed before computing the statistical model. This procedure allowed the identification of brain regions that were either very small or very large when compared to the rest of the sample and thus, might not be good representative of normality. For volumes and surfaces, this procedure was applied in proportion to eTIV (i.e. regional measure/eTIV). Since cortical thickness is not affected by eTIV, the outliers screening procedure was applied directly on the raw values. The number of outliers was below 1% for all regions (mean ±sd of all atlases: 0.45% ±0.10%) except the right long insular gyrus and central sulcus of the insula white surface (1.1%) and pericallosal sulcus volume (1.1%) of the Destrieux atlas. Detailed results can be found in the supplementary material as csv files.

### Regression models and statistical analyses

For each brain region measure, the normative values were produced following two linear regression models. First, a Model 1 was conducted with image quality predictors (voxel size, surface defect holes and CNR) and eTIV. Then, Model 2 with age and sex was applied on the residuals of Model 1. In order to respect the normality of the residuals, surface holes and all ventricles variables, except the 4th (3^rd^, lateral, inferior lateral and the sum of all ventricles), were log transformed. For ventricles and white matter regions, CNR of the total brain gray matter was used while for the brainstem subregions and hippocampal subfields, CNR from the whole brainstem and whole hippocampus were used, respectively. Quadratic and cubic terms for age, CNR and surface holes were included. Since voxel size has a relatively limited variability (mean: 1.02, std: 0.24, range: 0.18-2.2), we chose not to include quadratic and cubic terms for this variable. We also included all interactions except for voxel size (Model 1: eTIV X surface holes, eTIV x CNR, CNR X surface holes; Model 2: age X sex). Feature selection was conducted with a 10-fold cross-validation[58] backward elimination procedure, retaining the model with the subset of predictors that produced the lowest predicted residual sum of squares. For each selected final model, the fit of the data was assessed using R^2^ coefficient of determination:

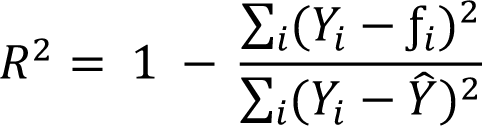

where the numerator is the residual sum of squares (*Y* is the value of the variable to be predicted and *f* is the predicted value), the denominator is the total sum of squares (9^<^ is the mean) and 4 is the index over subjects. To assess the unique contribution of each predictor, we used the lmg metric in the *R* package[59] relaimpo[60]. This metric is a R^2^ partitioned by averaging sequential sums of squares over all orderings of the predictors. Brain figures were made using the ggseg R package[61]. In order to compare the effects of each predictor, the sum of all relaimpo R^2^ terms related to each variable was computed (i.e. quadratic, cubic, and half of interaction values). For example, the variance explained by age includes the R^2^ sum for age, age^2^, age^3^, age X sex /2. When a term was not included within a model, its R^2^ value was given 0.

The models were verified by examining the difference between R^2^ of the training sample and R^2^ of the independent test sample of healthy controls. It was expected that the test R^2^ would be within 10% from the value of the training R^2^. Then, patterns of normality deviations were examined with the Z score effect sizes using the validation samples of healthy individuals and of individuals with AD and SZ.

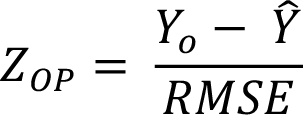

Z score effect sizes (Z_OP_) were obtained by subtracting the *Predicted value* (Ŷ) from the *Observed value* (*Y*_o_) divided by the root mean square error (RMSE) of the model predicting the value [62].

### Harmonization test

While the goal of NOMIS differs, we compared its results with twoharmonization procedures, NeuroCombat[16] and NeuroHarmonize[17] on the aseg volume and DKT cortical volume and thickness measures (matrix of 146 brain measures) from the harmonization dataset (SIMON, CIMA-Q and COMPASS-ND). We used the scanner identification number as “batch” (i.e. site) variable. For NeuroCombat, we also specified age and eTIV as covariates to preserve their effects. To compare harmonization procedures with NOMIS, after harmonization, eTIV was regressed out from the brain volume measures. Finally, to compare them on the same scale for statistical analyses on the variance and figure presentations, all measures were transformed into T and Z scores, respectively (see Supplementary Fig 2 as example).

We had three expectations following harmonization procedures. Compared to raw data, these procedures should:

- Reduce the variance of the measures from the 48 SIMON MRIs
- Maintain or increase effect sizes for hippocampi volumes and entorhinal thicknesses between CU, MCI, and AD groups
- Maintain or increase effect sizes for the relationships between hippocampi volumes and entorhinal thicknesses and episodic memory as measured by the delayed recall performance of the Rey Auditory Verbal Learning Test (RAVLT)[63] and Logical Memory Test[64].

The change in variance was assess using the quartile coefficient of dispersion (QCD):

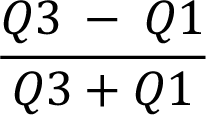

where Q1 and Q3 are the first and third quartile, respectively, and the Levene’s test for homogeneity of variance:

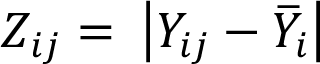

where *i* and *j* are the groups and the individuals, respectively. The Levene’s test is equivalent to a one-way between-groups analysis of variance (ANOVA) with the dependent variable being the absolute value of the difference between a score (Υ) and the mean of the group (Υ^-^). For each harmonization procedure, a one-way ANOVA on the QCD of the 146 measures before (raw) and after harmonization was conducted and a Levene’s test on each of the 146 measures was conducted.

All statistics were conducted using python’s module Scikit-learn[65] and Statsmodel[66].

## Results

As examples, figures in this report display results for subcortical volumes and DKT neocortical atlases volumes and thicknesses. Full detailed results for all atlases are provided as supplementary information as csv files.

### Model 1 – Image quality and eTIV

The R^2^ for model 1 ranged between 0.003 to 0.75, with a mean ±sd of 0.23 ± 0.15. The highest R^2^ were observed in brain measures with the largest volumes and surface areas (i.e. left and right white surface areas 0.82 and 0.81, and left and right pial surface areas 0.79 and 0.78, brain segmentation volume 0.75, supratentorial volume 0.75). Fig 2 shows the R^2^ portion due to image quality and eTIV for neocortical volumes and thicknesses of the DKT atlas parcellation, as well as subcortical volumes. Image quality had a substantial impact with a mean ±sd of 0.08 ±0.09, 0.14 ±0.05 and 0.13 ±0.05, for subcortical, neocortical volume and thickness values, respectively. A high amount of variance due to eTIV was observed in subcortical and neocortical volumes 0.22 ±0.12 and 0.26 ±0.08, respectively while it had nearly impact on cortical thickness measures 0.01 ±0.01.

**Fig 2.**
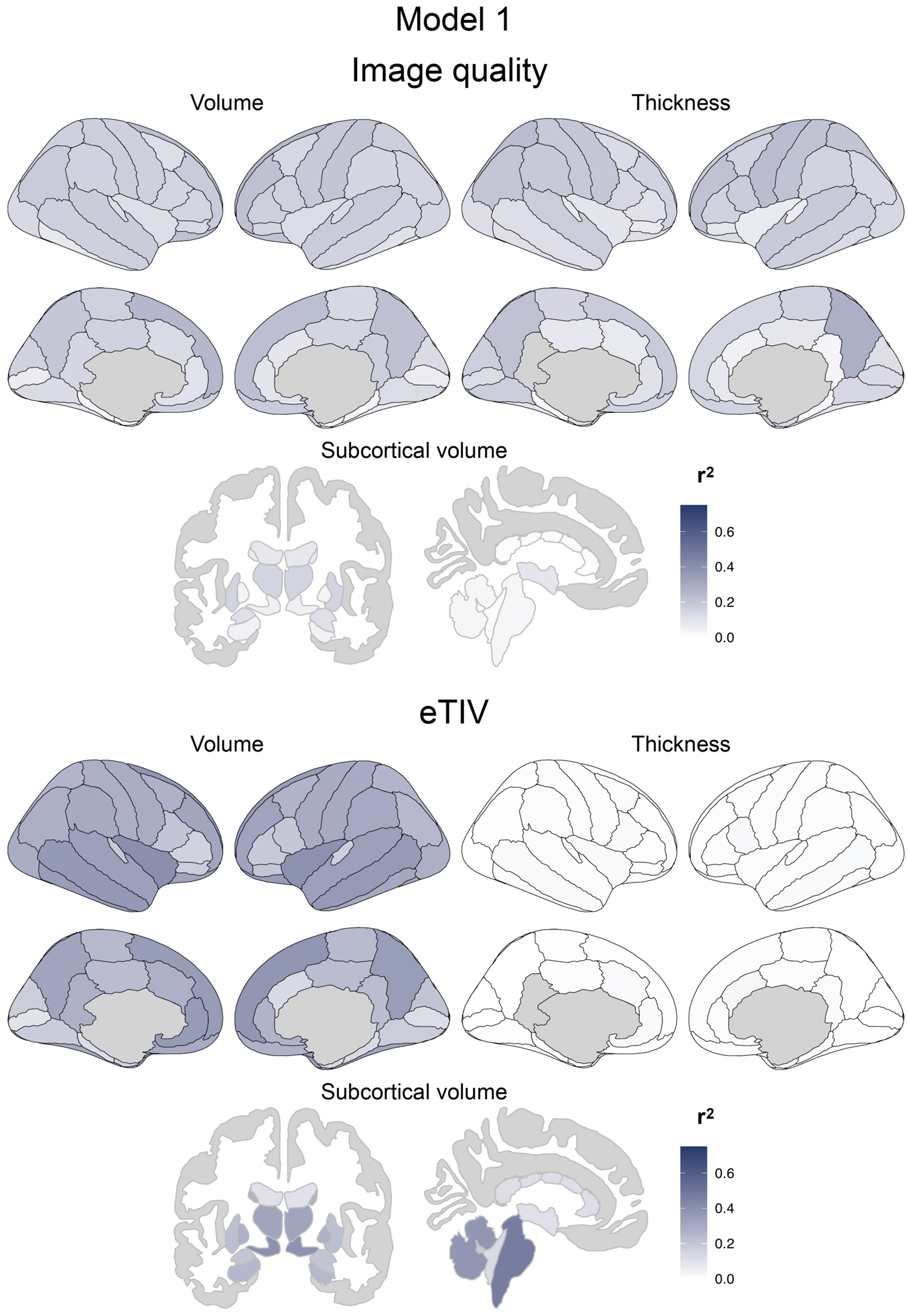
R^2^ from Model 1 for cortical volumes and thicknesses from the DKT atlas and subcortical volumes. Top: Variance due to image quality predictors. Bottom: Variance due to estimated intra-cranial volume (eTIV).

### Model 2 – Age and sex

The R^2^ for model 2 ranged between 0.02 to 0.51, with a mean ±sd of 0.23 ±0.14, 0.08 ±0.04 and 0.11 ±0.07, for subcortical volumes, neocortical volumes and thicknesses, respectively. One should note that the R^2^ in model 2 cannot be compared to that of model 1 since the total variance in model 2 is the remaining variance after model 1 (residuals). The highest R^2^ were observed in the largest regions and ventricles (i.e. all ventricles volume 0.51, brain segmentation volume 0.49, left and right lateral ventricles 0.49 and 0.49, supratentorial volumes 0.46). The lowest age and sex effects were generally on pial and white surface areas (0.00 ±0.10 and 0.03 ±0.02). Fig 3 illustrates the R^2^ for age and sex. As shown, sex did not explain much variance while age had a very different impact depending on the part of the brain with a higher impact in subcortical volumes and associative cortices, both in volume and thickness.

**Fig 3.**
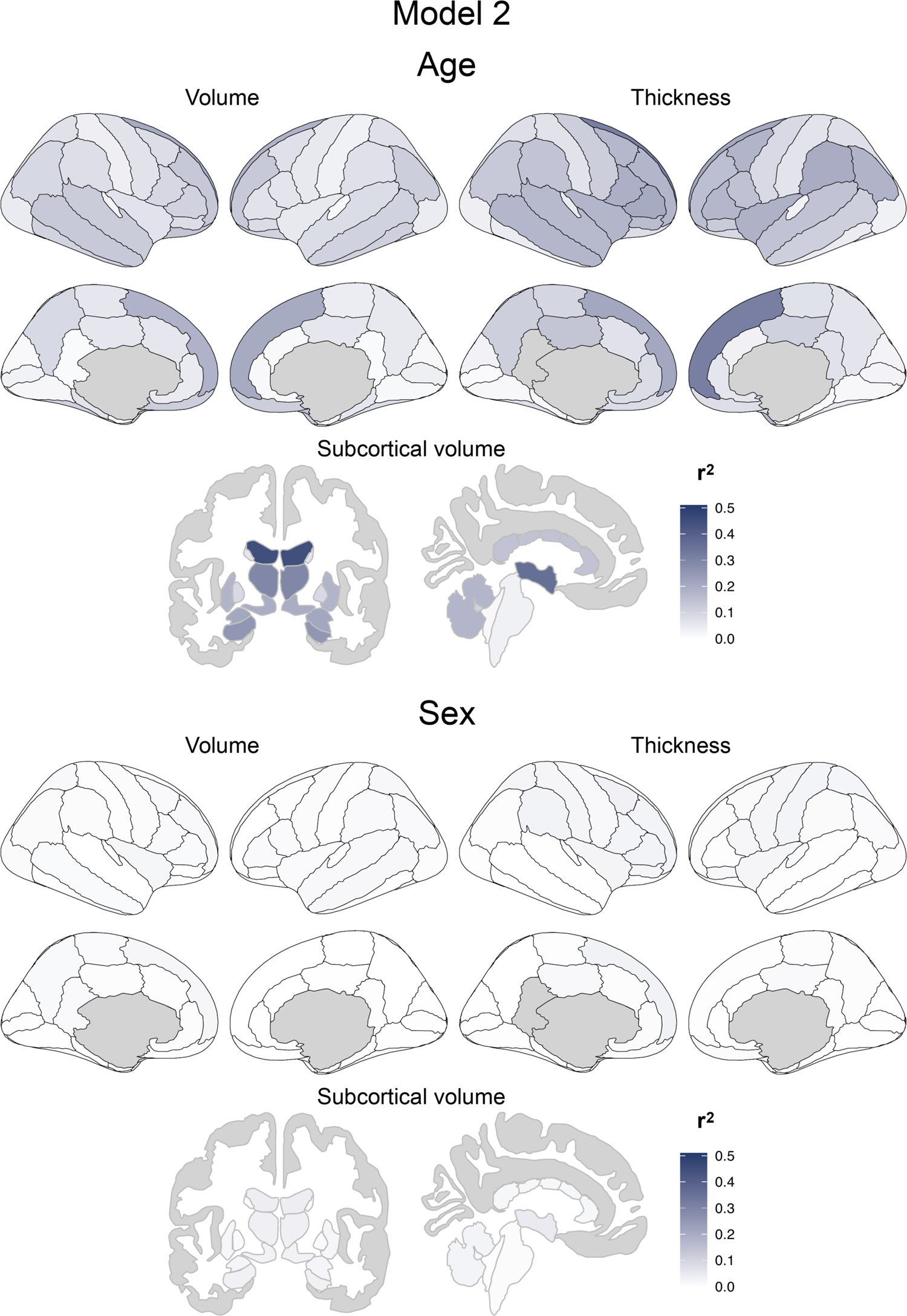
R^2^ from Model 2 for neocortical volumes and thicknesses from the DKT atlas and subcortical volumes. Top: Variance due to age. Bottom: Variance due to sex. One should note that the R^2^ in model 2 cannot be compared to that of model 1 since the total variance in model 2 is the remaining variance after model 1 (residuals).

### Models validation

Model 1 and Model 2 were examined in the independent test sample and nearly all models showed equivalent or higher R^2^ on the test set than on the training set (the difference test minus training for all atlases: Model 1 −0.026 ±0.027, Model 2 −0.005 ±0.018; Fig. 4). In model 1, the worse test differences were in the Destrieux atlas where 5 measures out of 592 were below −10%: pial surface areas of the left superior temporal sulcus (−0.13) and right fronto-marginal gyrus (of Wernicke) and sulcus (−0.12), the left subcallosal area, subcallosal gyrus volume (−0.10), the white surface area of the left lateral aspect of the superior temporal gyrus (0.10) and the right fronto-marginal gyrus (of Wernicke) and sulcus volume (−0.10). In Model 2, all measures had R^2^ differences below 10%, the worse being the left and right putamen volumes (−0.09 and −0.08). One should note that the models for these measures appear to be slightly less generalizable than the others.

**Fig 4.**
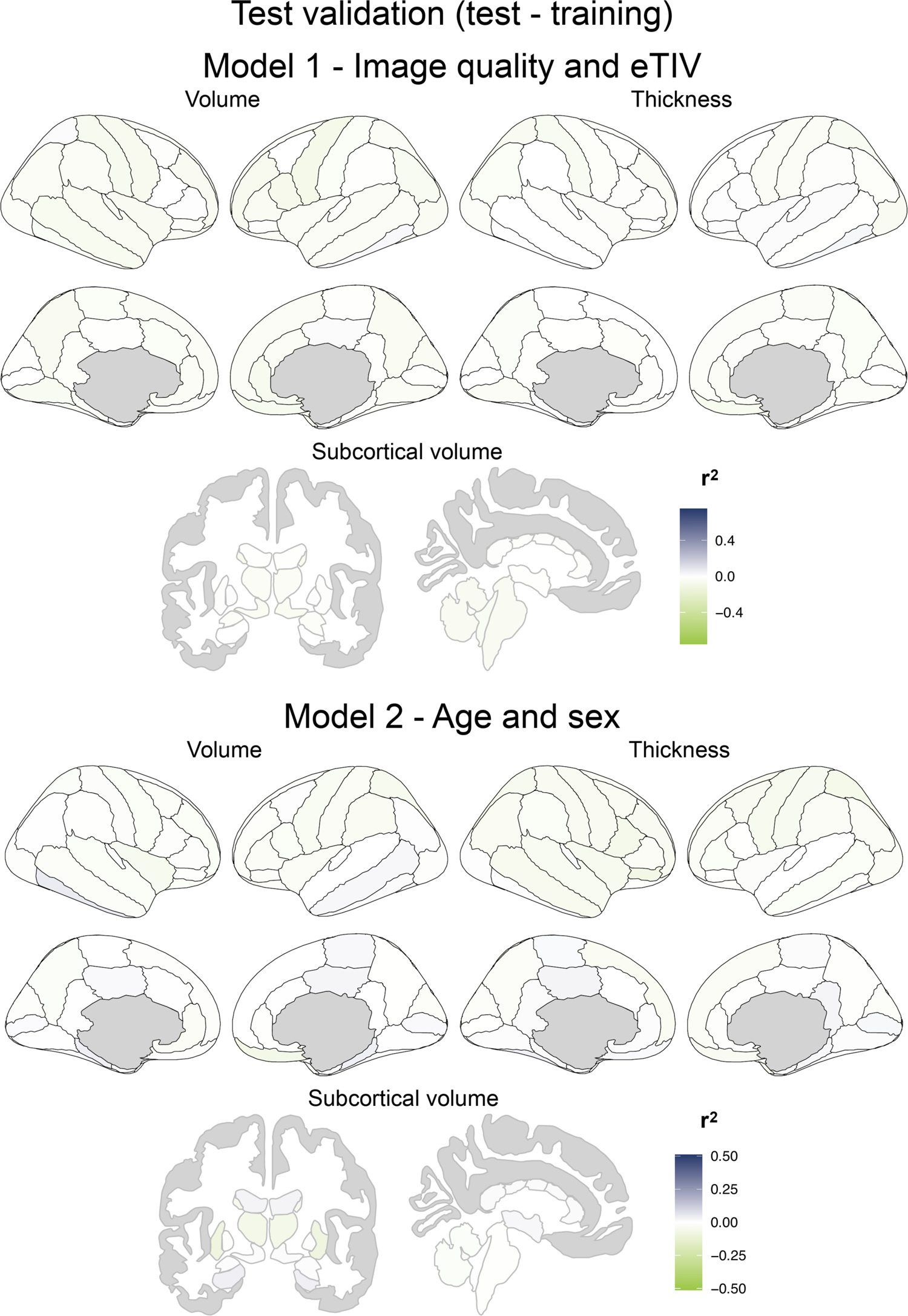
Difference of R^2^ between training and test samples. Top: Model 1 for Image quality and eTIV. Bottom: Model 2 for Age and sex.

Fig 5 and Fig 6 show the mean and std Z scores adjusted for age and sex when the models are applied on the independent young and older healthy controls samples. As expected, the mean Z scores were very close to 0 while the std were very close to 1 (mean ±std, Young, subcortical regions: −0.06 ±1.04, cortical volume: 0.04 ±1.13, cortical thickness: 0.04 ±0.99; Older, subcortical regions: 0.02 ±1.04, cortical volume: 0.01 ±0.96, cortical thickness: 0.01 ±1.02).

**Fig 5.**
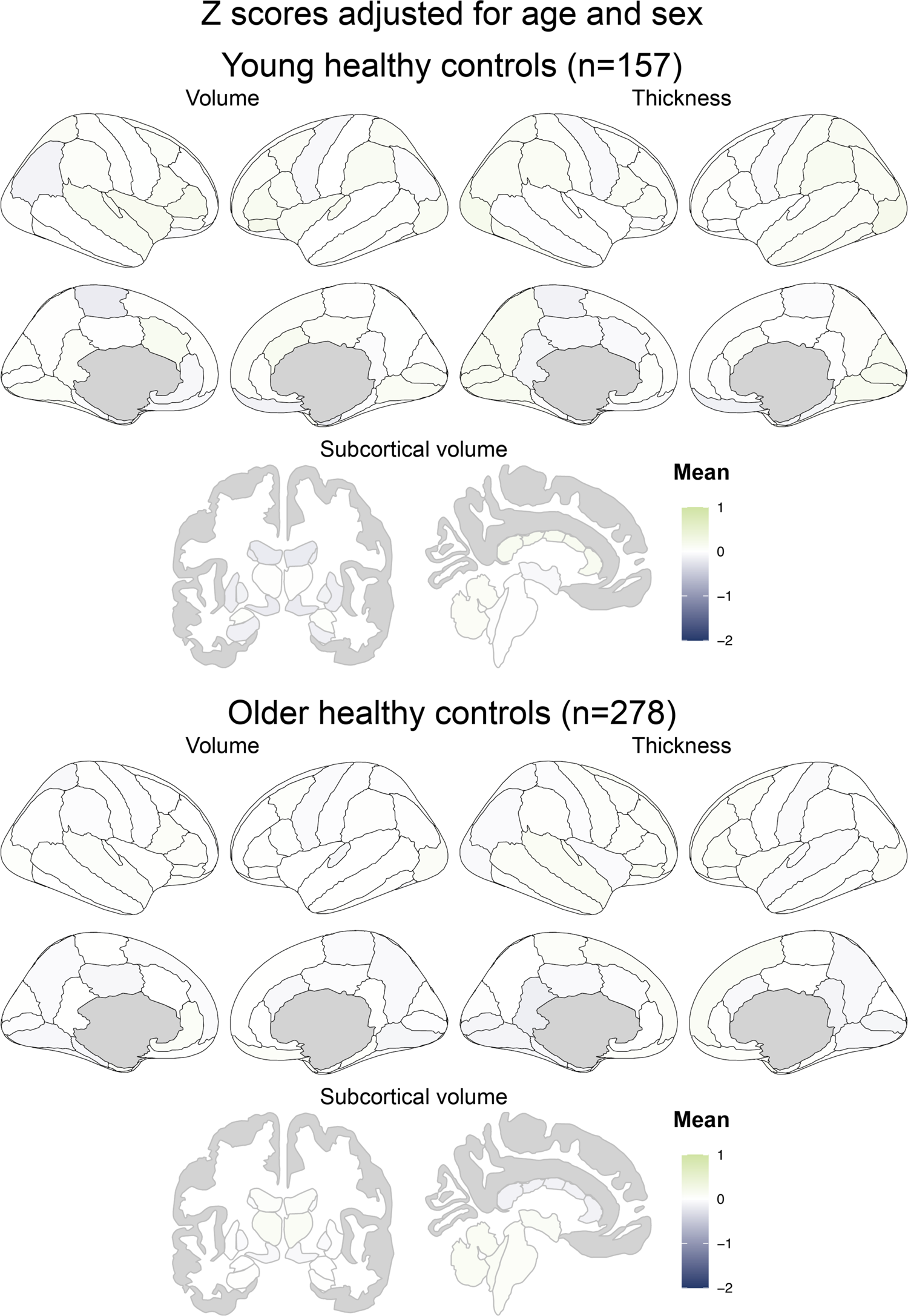
Mean normative Z scores on cortical volumes and thicknesses from the DKT atlas and subcortical volumes of young (18-34 years old) and older (65-92 years old) healthy participants. Note that the scaling chosen was to be comparable to that of Fig 9.

**Fig 6.**
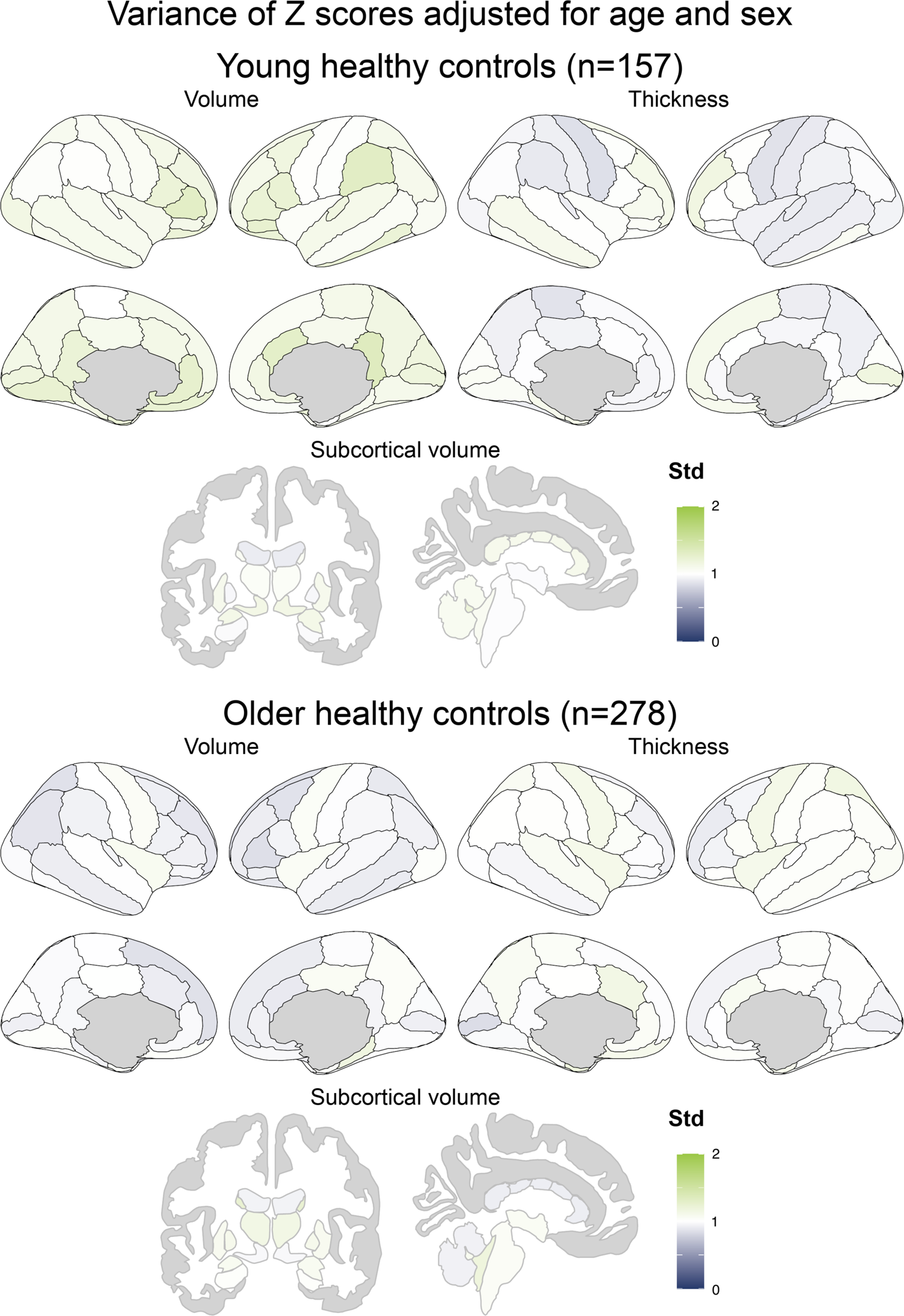
Variance of the normative Z scores on cortical volumes and thicknesses from the DKT atlas and subcortical volumes of young (18-34 years old) and older (65-92 years old) healthy participants. Std: standard deviation. The Std is expected to be near 1.

Using the independent healthy control test sample, Fig 7 and Fig 8 display examples of how the normative values remove the different effects on the left cortical thickness and volumes.

**Fig 7.**
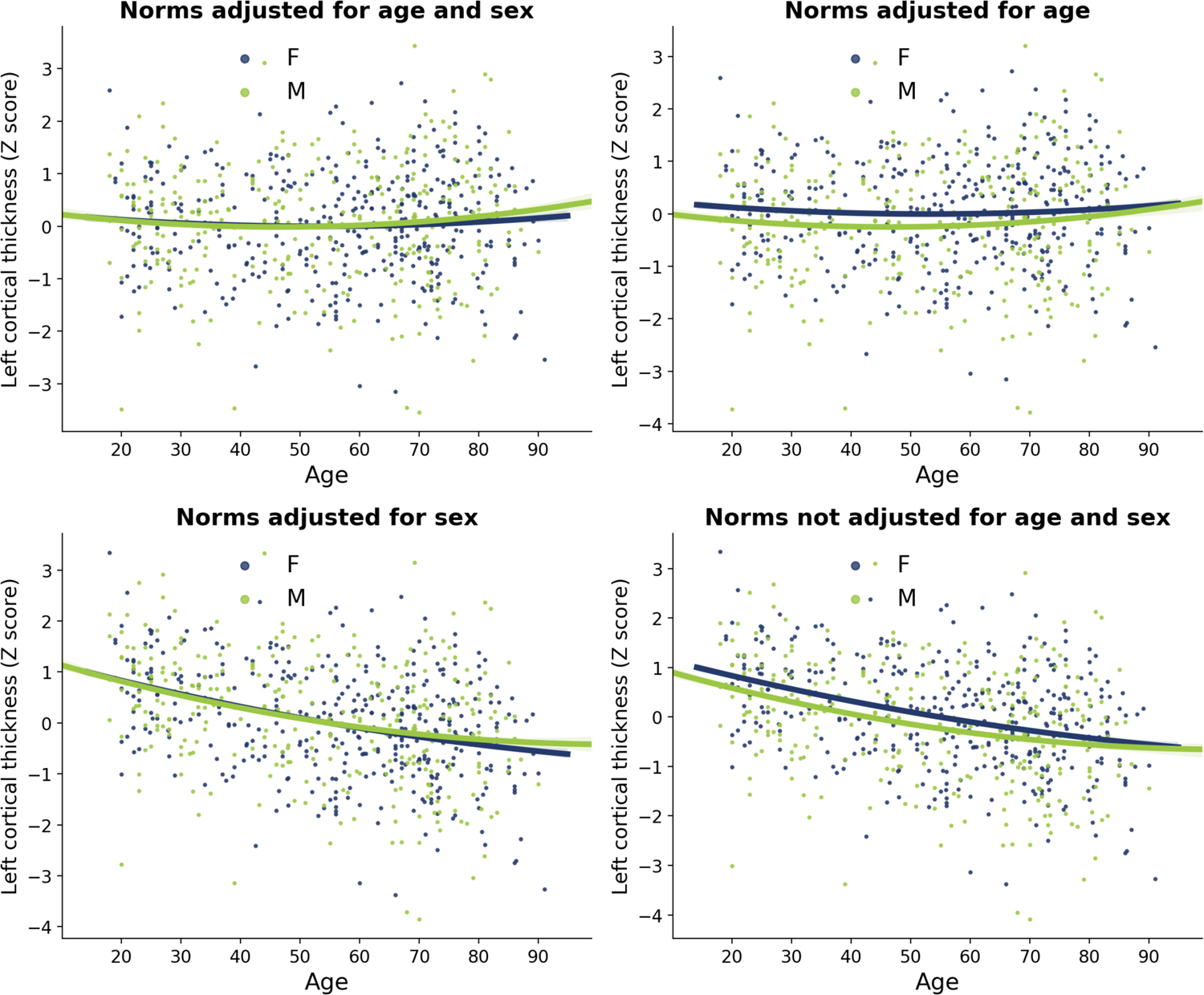
Example of the four NOMIS Z scores alternatives on the left cortical thickness values of the test sample. Note that all four alternative are adjusted for image quality.

**Fig 8.**
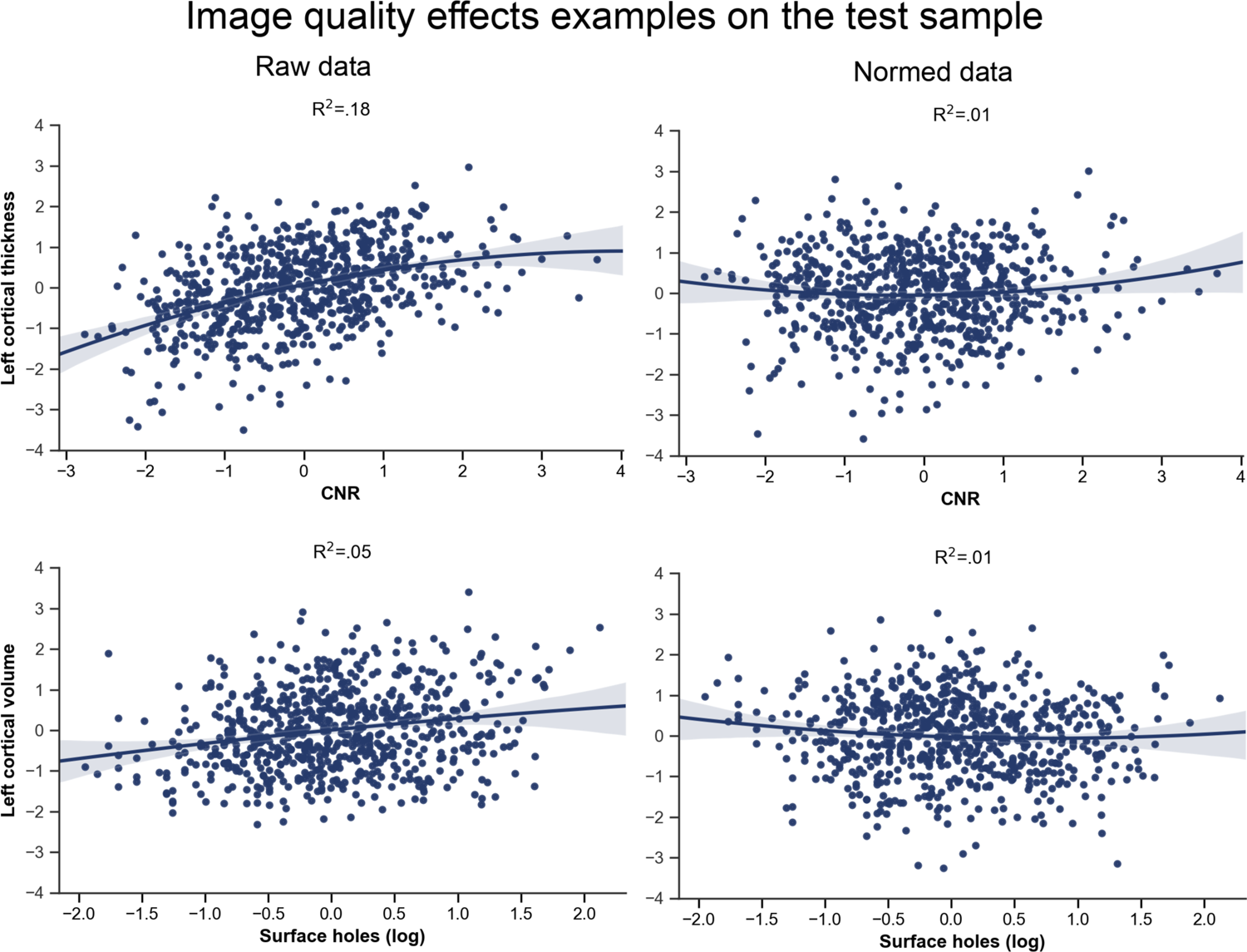
Examples of the impact of contrast-to-noise ratio (CNR) and surface holes on the raw and normed data of the test sample. Top: CNR on Left cortical thickness. Bottom: Surface holes on left cortical volume. Left: Raw data. Right: Normed data.

### Clinical validation

We validated the normative values in individuals with clinically ascertained Alzheimer’s disease and schizophrenia, which showed expected patterns of mean deviations from otherwise cognitively/behaviorally healthy individuals (Fig 9). In the Alzheimer’s disease group, the mean deviations from normality covered the frontal, temporal and parietal cortices with enlarged ventricles, but were especially more pronounced in the hippocampus and entorhinal cortex. In schizophrenia, atrophy was widespread to nearly all of the cortex. Supplementary Fig 3 displays the variance of the scores in those two groups.

**Fig 9.**
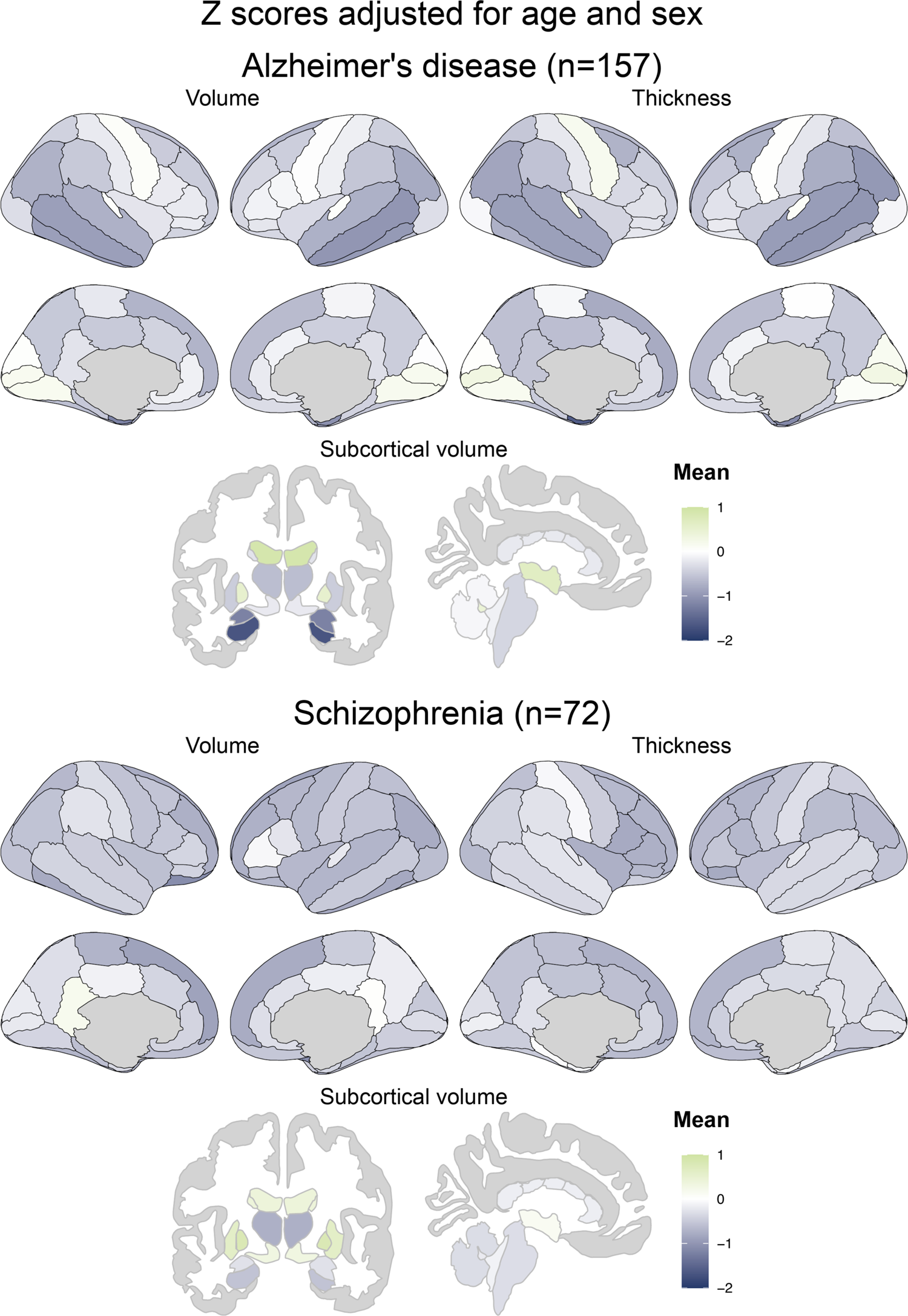
Mean normative Z scores on cortical volumes and thicknesses from the DKT atlas and subcortical volumes of participants with Alzheimer’s disease and with schizophrenia.

### Comparison of NOMIS and harmonization procedures

Fig 10 shows the SIMON subcortical and left cortical morphometric values (see Supplementary Fig 4 for right cortical values) before (raw) and after harmonization procedures and NOMIS normalization. Qualitatively, the variance of all measures was high before and after harmonization, as well as after NOMIS normalization. Fig 11 displays the QCD for subcortical and left cortical morphometric measures (see Supplementary Fig 5 for right cortical QCD values). QCD was highly different from a measure to another and globally harmonization procedures did not significantly lower QCD (NeuroCombat *F*: 1.96, *p*=.16; NeuroHarmonize *F*: 2.34, *p*=.13). On the other hand, NOMIS significantly reduced the QCD (*F*: 4.14, *p* = .04). Levene’s tests also indicated that the harmonization procedures and NOMIS values had equivalent variance than that of the raw values, even without correction for multiple comparison (*p* values ranging from .26 to .99). Fig 12 shows two examples of measures (left hippocampal volume and left entorhinal thickness) across the 12 different sites and reveals that, before and after harmonization or NOMIS, there is a high variability between sites, but also within some sites.

**Fig 10.**
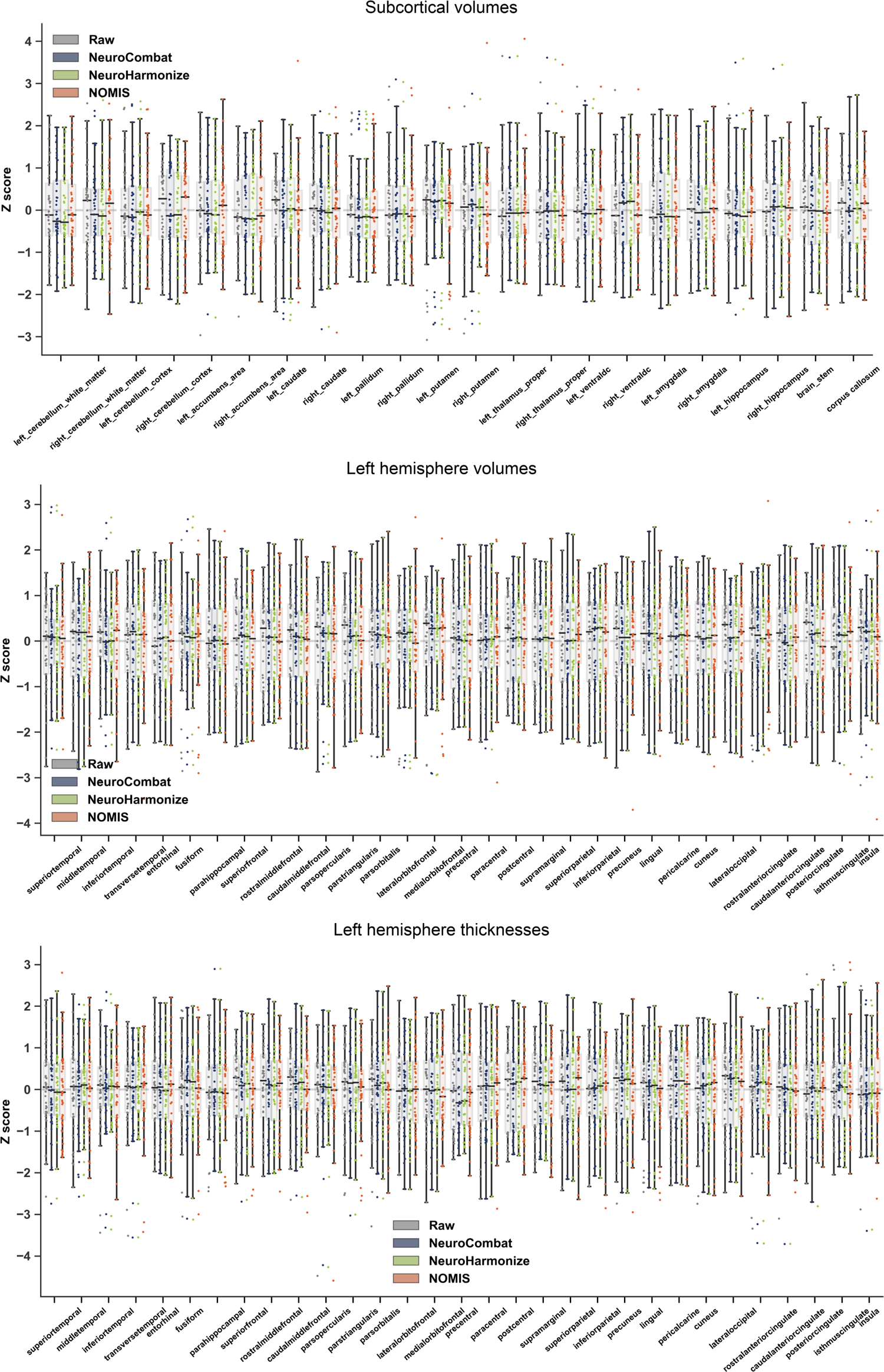
Boxplots showing the SIMON subcortical volumes and left neocortical volumes and thicknesses before (Raw) and after harmonization procedures (NeuroCombat and NeuroHarmonize) and NOMIS. Boxes show the first and third quartiles with the line denoting the median. Whiskers represent the lowest/highest datum still within 1.5 interquartile range (IQR) of the lower/higher quartile. Right neocortical volumes and thicknesses are shown in Supplementary Fig 4.

**Fig 11.**
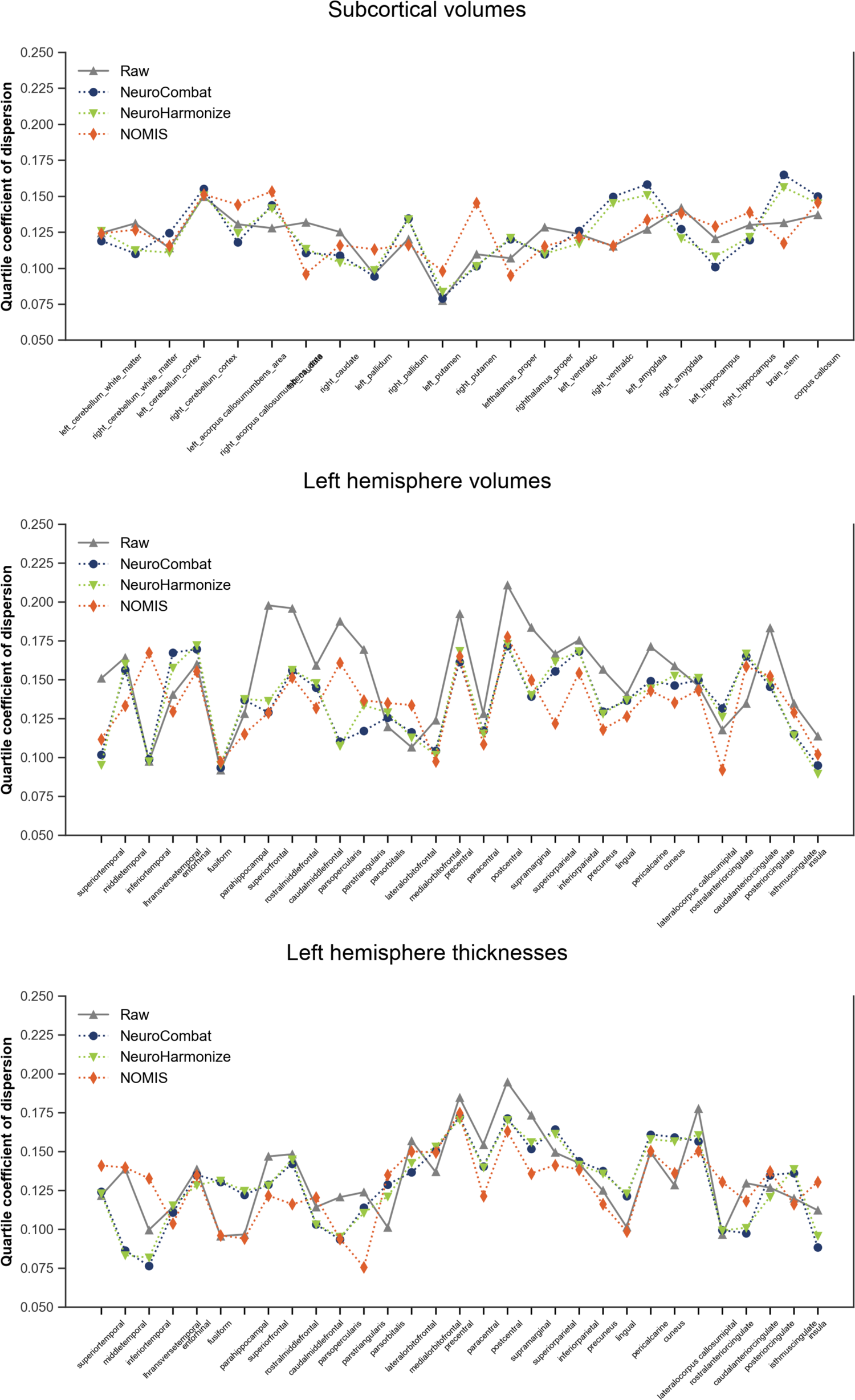
Quartile coefficient of dispersion of the SIMON subcortical volumes and left neocortical volumes and thicknesses before (Raw) and after harmonization procedures (NeuroCombat and NeuroHarmonize) and NOMIS. Values for the right neocortical volumes and thicknesses are shown in Supplementary Fig 5.

**Fig 12.**
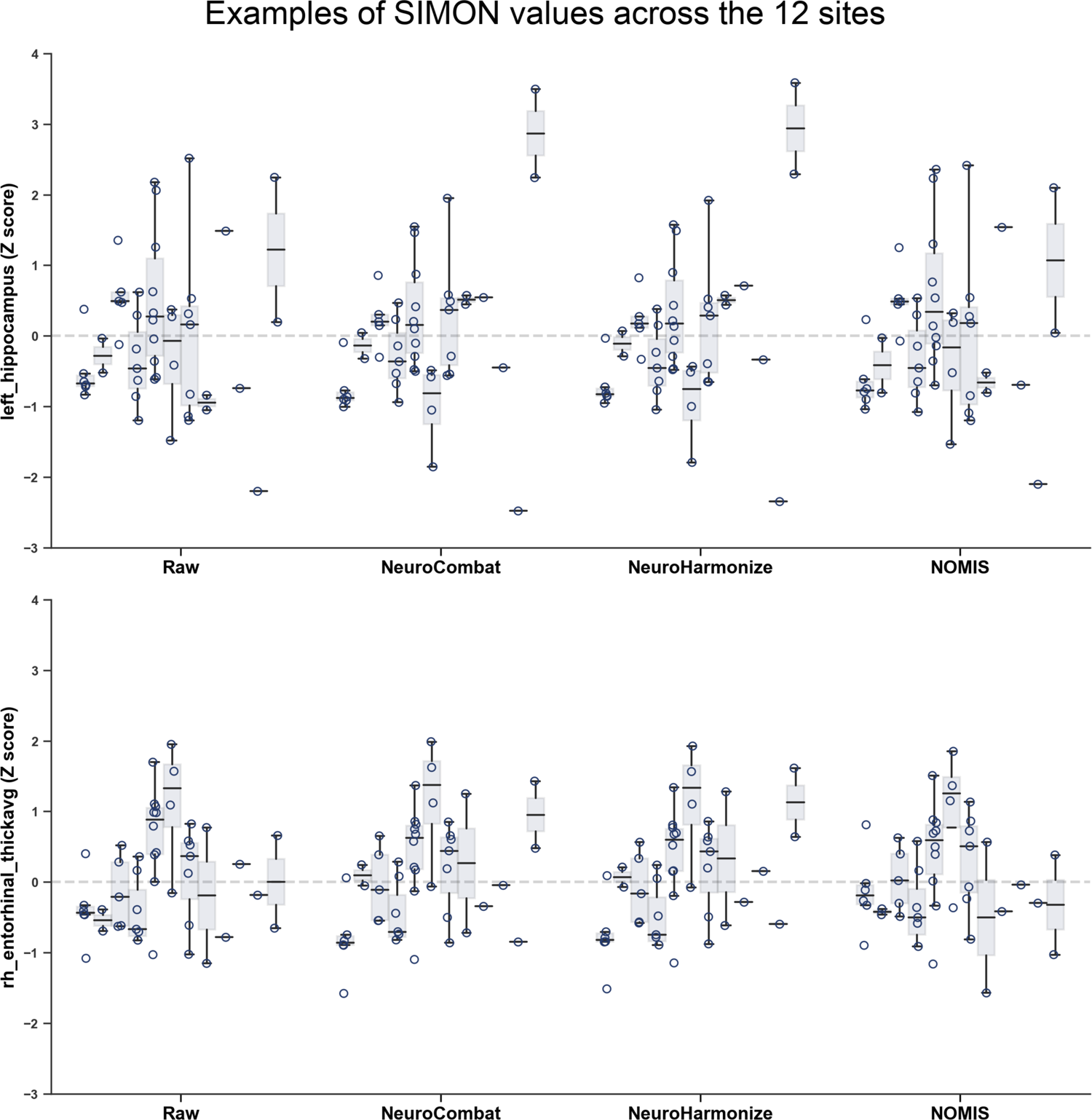
Boxplots showing the SIMON left hippocampal volumes and right entorhinal thicknesses before (Raw) and after harmonization procedures (NeuroCombat and NeuroHarmonize) and NOMIS for the 12 different sites. Boxes show the first and third quartiles with the line denoting the median. Whiskers represent the lowest/highest datum still within 1.5 interquartile range (IQR) of the lower/higher quartile.

Both harmonization procedures systematically diminished all effect sizes between CU and MCI (range: −0.01 to −0.11) and CU and AD (range: −0.09 to −0.20) groups. After NOMIS normalization, the effects sizes increased for hippocampal volumes (CU-MCI left: +0.14, right: +0.14; CU-AD: left: +0.12, right: +0.11) while it decreased for entorhinal volume (CU-MCI left: −0.01, right: −0.02; CU-AD: left: −0.08, right: −0.09) and thickness (CU-MCI left: −0.12, right: −0.16; CU-AD: left: −0.21, right: −0.28). These results are shown in Fig 13 (left hemisphere) and Supplementary Fig 6 (right hemisphere). Finally, we observed (Fig. 14) that both harmonization procedures systematically lowered the magnitude of the correlations between the six morphometric values and episodic memory scores. NOMIS on the other hand increased the correlations with hippocampal volumes, slightly decreased the ones with entorhinal cortices and performed similarly to harmonization procedures for entorhinal cortices thicknesses.

**Fig 13.**
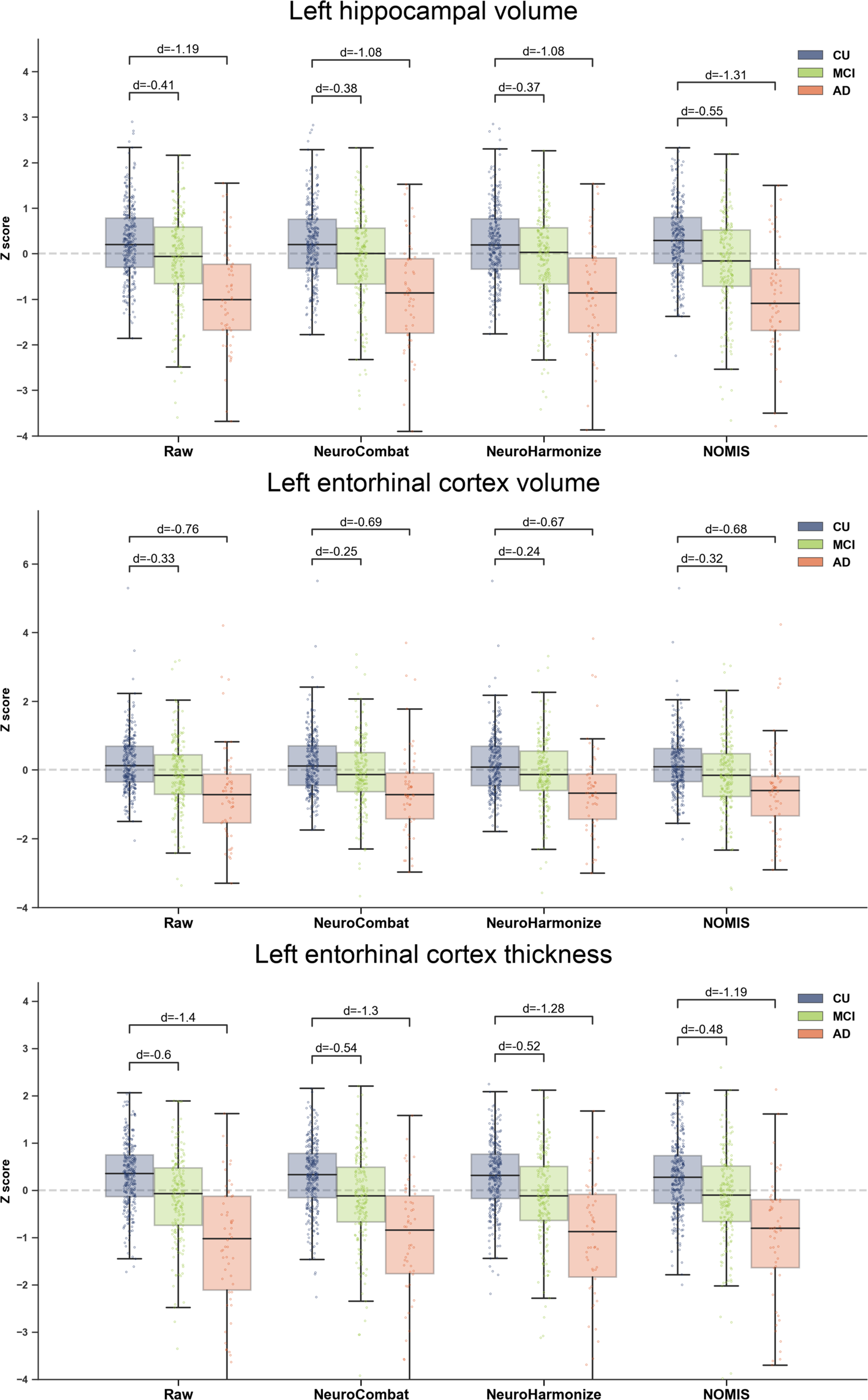
Boxplots showing the effect sizes (Cohen’s d) between cognitively unimpaired (CU), mild cognitive impairment (MCI) and Alzheimer’s disease (AD) groups for the left hippocampal volume and left entorhinal volume and thickness before (Raw) and after harmonization procedures (NeuroCombat and NeuroHarmonize) and NOMIS. Boxes show the first and third quartiles with the line denoting the median. Whiskers represent the lowest/highest datum still within 1.5 interquartile range (IQR) of the lower/higher quartile. Values for the right hemisphere are shown in Supplementary Fig 6.

**Fig 14.**
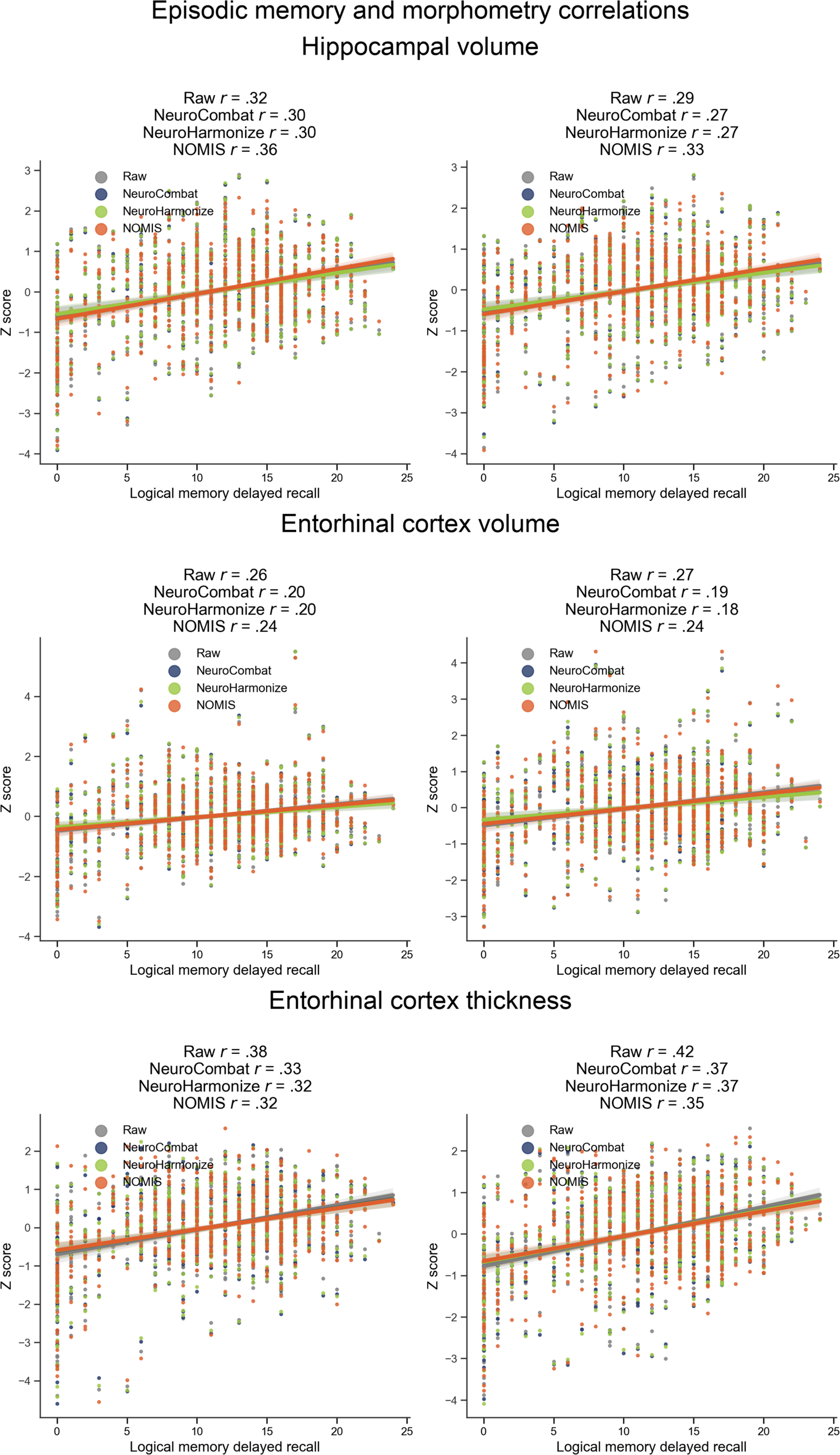
Correlations between episodic memory score and morphometric measures before (Raw) and after harmonization procedures (NeuroCombat and NeuroHarmonize) and NOMIS.

## Discussion

Recent initiatives for morphometric normative data includes percentile fitting curves on subcortical regions [67], deep learning-based segmentation of subcortical regions and cortical lobes for east Asians [68], and yearly percentage of brain volume changes [69]. To our knowledge, there is no other automated calculator for normative morphometric values available to researchers except the one from our previous work using *FreeSurfer* 5.3 (https://github.com/medicslab/mNormsFS53). These prior normative data from our group[5–7] were relatively limited in terms of atlases and sample size. With nearly seven thousand participants and 1,344 brain measures, NOMIS offers a comprehensive neuromorphometric normative tool based on a very large sample. In addition, an innovation of NOMIS is its flexibility.

Depending on the user need, it has four versions of Z-score adjusted on different sets of variables. All versions include head size and image quality, but can also take into account age and/or sex or without age and sex. Therefore, research groups looking for traditional norms, as well as others wanting to lower the variance due to head size and image quality while preserving age and/or sex variances can take advantage of NOMIS. Another strength of NOMIS is that the normative values were created on a various amalgam of cognitively healthy participants from multiple countries, with data acquired from a wide variety of MRI scanners and image quality, maximizing its generalizability. A novelty to prior existing normative data, is the addition of the image quality impact on the morphometry measures. Figures 2 shows that its effect is not trivial on cortical volume and thickness. As shown by our results, our new normative data should help to remove some undesirable variance due to scanners and image quality. Furthermore, the results from NOMIS also show that in independent samples, the Z scores behaved as expected, that is with a mean of 0 and standard deviation of 1 in healthy individuals and with marked mean deviations targeted to the medial temporal lobes in participants with AD and throughout the cortex in participants with SZ.

Despite these strengths, users should keep in mind that before using NOMIS, it is mandatory to verify *FreeSurfer* segmentations and that while it will remove parts of variance due to head size and image quality, it won’t correct for segmentation errors or image artefacts. Moreover, the normative sample, comprised essentially of research volunteers in academic-led environments, was recruited using a non-probability sampling method and may not be representative of the targeted population by the user.

### Norms and multi-site data harmonization

The main aim of the normative values is to quantify the deviation from normality of measurements for a new individual (i.e. one who is not in the sample used to define normality). Because the norms remove some variance due to image quality, they can also be a useful and simple way of lowering the noise between scanners in multi-site studies. However, norms should not be considered an optimal technique to remove variances in multi-site studies; other strategies are meant to specifically tackle multi-site variance. Studies should use in fact a combination of approaches, including harmonized procedures for data acquisition, normative values such as the one proposed herein, and post-hoc correction.

A harmonized scanning procedure, such as the Canadian Dementia Imaging Protocol (CDIP)[70], addresses variations due to parameter and sequence dissimilarities, including quality control and assurance, for example through scanning at all sites of an object of known geometric and contrast properties (i.e. a “phantom”) as well as human volunteers. Recent data from the SIMON dataset [21], including non-harmonized and CDIP-harmonized scans, demonstrated how using a harmonized protocol reduces variability across sites; however, some notable variance remained[71, 72], (see also Figs 10-12). The idea of a harmonized protocol is however limited to specific initiatives due to the high amount of resources it requires to implement.

Post-hoc harmonization procedures on the other hand have been developed to pool data from different sites in large studies. Various of procedures have been proposed [16–19, 73] and aim to lower differences in morphometric data between sites by generally applying scaling corrections based on differences in the morphometric data themselves. The scaling corrections are applicable for the sites/scanners included in the analysis and not for future sites/scanners or data. This makes such post-hoc correction analysis-specific and needs to be conducted each time some data are removed or added to an analysis. Such an approach can be very useful for large multi-centric studies but is not applicable for generating normative values aiming to be applied on future data. It is also vulnerable to selection bias since the scaling factors are not based on the images or scanner characteristics, but on the difference of data between sites/scanners[18, 19]. Thus, distinct characteristics of the participants at a given site can affect the scaling factors and post-hoc scaling factors should be used when the aim of a study is not vulnerable to sources of variance between sites that are not related to image acquisition.

We compared NOMIS values to two post-hoc harmonization procedures, namely NeuroCombat[16] and NeuroHarmonize[17] and while globally NOMIS slightly lowered the variance of the values from the same individuals originating from 12 different scanners, these two procedures were worse than NOMIS and did not significantly reduce true variance induced by different scanners. We also verified effect sizes of well-established effects in MCI and AD participants and once again the harmonization procedures were either similar or worse than NOMIS. NeuroCombat and NeuroHarmonize systematically lowered the morphometric differences between CU, MCI and AD participants while NOMIS lowered the entorhinal volume and thickness effect sizes and increased the hippocampal volume differences between these groups. These results suggest that caution should be exercised when using post-hoc harmonization; the use of a calibration technique (e.g. repeated scans of human volunteers as part of the study) is strongly encouraged.

### Using NOMIS

The NOMIS tool is a user-friendly automated script in Python, freely accessible (https://github.com/medicslab/NOMIS). Users only need to pre-process their images with *FreeSurfer* 6.0 using automated directive parameters, then specify the individuals’ characteristics to the script, which will automatically compute Z-scores based on the *FreeSurfer* output. One can choose the version of the Z-score by including in the csv file only the variables that need to be adjusted and the script automatically selects the appropriate version of predictors. The predictive models and all statistical parameters are provided along with the script. We anticipate that this tool will be of broad interest to the neuroscientific community.

## Acknowledgments

This study comprises multiple samples of healthy individuals. We wish to thank all principal investigators who collected these datasets and agreed to let them accessible.

Autism Brain Imaging Data Exchange (ABIDE): Primary support for the work by Adriana Di Martino was provided by the NIMH (K23MH087770) and the Leon Levy Foundation. Primary support for the work by Michael P. Milham and the INDI team was provided by gifts from Joseph P. Healy and the Stavros Niarchos Foundation to the Child Mind Institute, as well as by an NIMH award to MPM (R03MH096321). http://fcon_1000.projects.nitrc.org/indi/abide/ Alzheimer’s Disease Neuroimaging Initiative (ADNI): The investigators within the ADNI contributed to the design and implementation of ADNI and/or provided data but did not participate in analysis or writing of this report. A complete listing of ADNI investigators can be found at: http://adni.loni.usc.edu/wp-content/uploads/how_to_apply/ADNI_Acknowledgement_List.pdf. ADNI was funded by the Alzheimer’s Disease Neuroimaging Initiative (ADNI) (National Institutes of Health Grant U01 AG024904) and DOD ADNI (Department of Defense award number W81XWH-12-2-0012). ADNI is funded by the National Institute on Aging, the National Institute of Biomedical Imaging and Bioengineering, and through generous contributions from the following: AbbVie, Alzheimer’s Association; Alzheimer’s Drug Discovery Foundation; Araclon Biotech; BioClinica, Inc.; Biogen; Bristol-Myers Squibb Company; CereSpir, Inc.; Cogstate; Eisai Inc.; Elan Pharmaceuticals, Inc.; Eli Lilly and Company; EuroImmun; F. Hoffmann-La Roche Ltd and its affiliated company Genentech, Inc.; Fujirebio; GE Healthcare; IXICO Ltd.;Janssen Alzheimer Immunotherapy Research & Development, LLC.; Johnson & Johnson Pharmaceutical Research & Development LLC.; Lumosity; Lundbeck; Merck & Co., Inc.; Meso Scale Diagnostics, LLC.; NeuroRx Research; Neurotrack Technologies; Novartis Pharmaceuticals Corporation; Pfizer Inc.; Piramal Imaging; Servier; Takeda Pharmaceutical Company; and Transition Therapeutics. The Canadian Institutes of Health Research is providing funds to support ADNI clinical sites in Canada. Private sector contributions are facilitated by the Foundation for the National Institutes of Health (www.fnih.org). The grantee organization is the Northern California Institute for Research and Education, and the study is coordinated by the Alzheimer’s Therapeutic Research Institute at the University of Southern California. ADNI data are disseminated by the Laboratory for Neuro Imaging at the University of Southern California. http://adni.loni.usc.edu/

Australian Imaging Biomarkers and Lifestyle flagship study of ageing (AIBL): Part of the data used in this study was obtained from the Australian Imaging Biomarkers and Lifestyle flagship study of ageing (AIBL) funded by the Commonwealth Scientific and Industrial Research Organisation (CSIRO) which was made available at the ADNI database (www.loni.usc.edu/ADNI). The AIBL researchers contributed data but did not participate in analysis or writing of this report. AIBL researchers are listed at www.aibl.csiro.au

Berlin Mind and Brain (Margulies, Villringer) CoRR sample (BMB). Zuo, X.N., et al. (2014). An open science resource for establishing reliability and reproducibility in functional connectomics. *Scientific data, 1*, 140049. doi: 10.1038/sdata.2014.49. http://fcon_1000.projects.nitrc.org/indi/CoRR/html/bmb_1.html

Cambridge Centre for Ageing and Neuroscience (CamCAN): CamCAN funding was provided by the UK Biotechnology and Biological Sciences Research Council (grant number BB/H008217/1), together with support from the UK Medical Research Council and University of Cambridge, UK. http://www.mrc-cbu.cam.ac.uk/datasets/camcan/

Center of Biomedical Research Excellence (COBRE): The imaging data and phenotypic information was collected and shared by the Mind Research Network and the University of New Mexico funded by a National Institute of Health COBRE: 1P20RR021938-01A2. http://fcon_1000.projects.nitrc.org/indi/retro/cobre.html

Cleveland Clinic (Cleveland CCF): Funded by the National Multiple Sclerosis Society. http://fcon_1000.projects.nitrc.org/indi/retro/ClevelandCCF.html

Comprehensive Assessment of Neurodegeneration and Dementia (COMPASS-ND) study: The COMPASS-ND study is conducted by the Canadian Consortium on Neurodegeneration in Aging (CCNA; www.ccna-ccnv.ca). The CCNA is supported by a grant from the Canadian Institutes of Health Research (CIHR) with funding from several partners.

Consortium for the Early Identification of Alzheimer’s Disease (CIMA-Q): Part of the data used in this article were obtained from the Consortium pour l’identification précoce de la maladie Alzheimer - Québec (CIMA-Q). As such, the investigators within the CIMA-Q contributed to the design, the implementation, the acquisition of clinical, cognitive, and neuroimaging data and biological samples. A list of the CIMA-Q investigators is available on <u>cima-q.ca</u>. CIMA-Q was funded in 2013 with a $2,500,000 grant from the Fonds d’Innovation Pfizer - Fond de Recherche Québec – Santé sur la maladie d’Alzheimer et les maladies apparentées.

Dallas Lifespan Brain Study (DLBS): This study is supported by the Center for Vital Longevity, the University of Texas at Dallas, the University of Texas Southwestern Medical Center, the National Institutes of Health and Aging, AVID Radiopharmaceuticals, the Aging Mind Foundation and the Alzheimer’s Association. http://fcon_1000.projects.nitrc.org/indi/retro/dlbs.html

FIND lab sample. Funded by the Dana Foundation; John Douglas French Alzheimer’s Foundation; National Institutes of Health (AT005733, HD059205,HD057610, NS073498, NS058899). http://fcon_1000.projects.nitrc.org/indi/retro/find_stanford.html

Functional Biomedical Informatics Research Network (FBIRN): Provided by the Biomedical Informatics Research Network under the following support: U24-RR021992, by the National Center for Research Resources at the National Institutes of Health, U.S.A. http://www.birncommunity.org/resources/data/

Lifespan Human Connectome Project in Aging (HCP-Aging): HCP-Aging data were obtained from the National Institute of Mental Health (NIMH) Data Archive (NDA). NDA is a collaborative informatics system created by the National Institutes of Health to provide a national resource to support and accelerate research in mental health. Dataset identifier: http://dx.doi.org/10.15154/1520138. This manuscript reflects the views of the authors and may not reflect the opinions or views of the NIH or of the Submitters submitting original data to NDA. http://nda.nih.gov

International Consortium for Brain Mapping (ICBM). The ICBM (Principal Investigator: John Mazziotta, MD, PhD) was funded was provided by the National Institute of Biomedical Imaging and BioEngineering. ICBM is the result of efforts of co-investigators from UCLA, Montreal Neurologic Institute, University of Texas at San Antonio, and the Institute of Medicine, Juelich/Heinrich Heine University - Germany.” https://ida.loni.usc.edu/login.jsp?project=ICBM

Information eXtraction from Images (IXI): Data collected as part of the project EPSRC GR/S21533/02 - http://brain-development.org/ixi-dataset/

F.M. Kirby Research Center neuroimaging reproducibility data (KIRBY-21). Landman, B.A. et al. “Multi-Parametric Neuroimaging Reproducibility: A 3T Resource Study”, NeuroImage. (2010) NIHMS/PMC:252138 doi:10.1016/j.neuroimage.2010.11.047 https://www.nitrc.org/projects/multimodal

Minimal Interval Resonance Imaging in Alzheimer’s Disease (MIRIAD): The MIRIAD investigators did not participate in analysis or writing of this report. The MIRIAD dataset is made available through the support of the UK Alzheimer’s Society (RF116). The original data collection was funded through an unrestricted educational grant from GlaxoSmithKline (6GKC). http://miriad.drc.ion.ucl.ac.uk

National Alzheimer’s Coordinating Center (NACC): The NACC database is funded by NIA/NIH Grant U01 AG016976. NACC data are contributed by the NIA-funded ADCs: P30 AG019610 (PI Eric Reiman, MD), P30 AG013846 (PI Neil Kowall, MD), P30 AG062428-01 (PI James Leverenz, MD) P50 AG008702 (PI Scott Small, MD), P50 AG025688 (PI Allan Levey, MD, PhD), P50 AG047266 (PI Todd Golde, MD, PhD), P30 AG010133 (PI Andrew Saykin, PsyD), P50 AG005146 (PI Marilyn Albert, PhD), P30 AG062421-01 (PI Bradley Hyman, MD, PhD), P30 AG062422-01 (PI Ronald Petersen, MD, PhD), P50 AG005138 (PI Mary Sano, PhD), P30 AG008051 (PI Thomas Wisniewski, MD), P30 AG013854 (PI Robert Vassar, PhD), P30 AG008017 (PI Jeffrey Kaye, MD), P30 AG010161 (PI David Bennett, MD), P50 AG047366 (PI Victor Henderson, MD, MS), P30 AG010129 (PI Charles DeCarli, MD), P50 AG016573 (PI Frank LaFerla, PhD), P30 AG062429-01(PI James Brewer, MD, PhD), P50 AG023501 (PI Bruce Miller, MD), P30 AG035982 (PI Russell Swerdlow, MD), P30 AG028383 (PI Linda Van Eldik, PhD), P30 AG053760 (PI Henry Paulson, MD, PhD), P30 AG010124 (PI John Trojanowski, MD, PhD), P50 AG005133 (PI Oscar Lopez, MD), P50 AG005142 (PI Helena Chui, MD), P30 AG012300 (PI Roger Rosenberg, MD), P30 AG049638 (PI Suzanne Craft, PhD), P50 AG005136 (PI Thomas Grabowski, MD), P30 AG062715-01 (PI Sanjay Asthana, MD, FRCP), P50 AG005681 (PI John Morris, MD), P50 AG047270 (PI Stephen Strittmatter, MD, PhD). https://www.alz.washington.edu/

National Database for Autism Research (NDAR): Data were obtained from the National Institute of Mental Health (NIMH) Data Archive (NDA). NDA is a collaborative informatics system created by the National Institutes of Health to provide a national resource to support and accelerate research in mental health. Dataset identifier: http://dx.doi.org/10.15154/1520138. This manuscript reflects the views of the authors and may not reflect the opinions or views of the NIH or of the Submitters submitting original data to NDA. http://nda.nih.gov

Nathan Kline Institute Rockland (NKI-R) sample (NKI-RS) and Enhanced Sample (NKI-RES): Principal support for the NKI-RES project is provided by the NIMH BRAINS R01MH094639-01. Funding for key personnel also provided in part by the New York State Office of Mental Health and Research Foundation for Mental Hygiene. Funding for the decompression and augmentation of administrative and phenotypic protocols provided by a grant from the Child Mind Institute (1FDN2012-1). Additional personnel support provided by the Center for the Developing Brain at the Child Mind Institute, as well as NIMH R01MH081218, R01MH083246, and R21MH084126. Project support also provided by the NKI Center for Advanced Brain Imaging (CABI), the Brain Research Foundation, the Stavros Niarchos Foundation and the NIH P50 MH086385-S1 (NKI-RS). http://fcon_1000.projects.nitrc.org/indi/pro/nki.html http://fcon_1000.projects.nitrc.org/indi/enhanced/

Open access series of imaging studies (OASIS): The OASIS project was funded by grants P50 AG05681, P01 AG03991, R01 AG021910, P50 MH071616, U24 RR021382, and R01 MH56584. http://www.oasis-brains.org/

POWER: This database was supported by NIH R21NS061144 R01NS32979 R01HD057076 U54MH091657 K23DC006638 P50 MH71616 P60 DK020579-31, McDonnell Foundation Collaborative Action Award, NSF IGERT DGE-0548890, Simon’s Foundation Autism Research Initiative grant, Burroughs Wellcome Fund, Charles A. Dana Foundation, Brooks Family Fund, Tourette Syndrome Association, Barnes-Jewish Hospital Foundation, McDonnell Center for Systems Neuroscience, Alvin J. Siteman Cancer Center, American Hearing Research Foundation grant, Diabetes Research and Training Center at Washington University grant. http://fcon_1000.projects.nitrc.org/indi/retro/Power2012.html

Parkinson’s Progression Markers Initiative (PPMI): PPMI – a public-private partnership – is funded by the Michael J. Fox Foundation for Parkinson’s Research and funding partners, including Abbvie, Allergan, Amathus, Avid Radiopharmaceuticals, Biogen Idec, BioLegend, Bristol-Myers, Celgene, Cenali, Covance, GE Healthcare, Genentech, GlaxoSmithKline, Glolub Capital, Handl Therapeutics, Insitro, Janssen Neuroscience, Eli Lilly and Company, Lundbeck, Merck, Meso Scale Discovery, Neurocrine, Pfizer, Piramal, Prevail, Roche, Sanofi Genzyme, Servier, Takeda, Teva, UCB, Verily, and Voyager Therapeutics. See http://www.ppmi-info.org for further details.

Southwest University Adult Lifespan Dataset (SALD): SALD was supported by the National Natural Science Foundation of China (31470981; 31571137; 31500885), National Outstanding young people plan, the Program for the Top Young Talents by Chongqing, the Fundamental Research Funds for the Central Universities (SWU1509383,SWU1509451,SWU1609177), Natural Science Foundation of Chongqing (cstc2015jcyjA10106), Fok Ying Tung Education Foundation (151023), General Financial Grant from the China Postdoctoral Science Foundation (2015M572423, 2015M580767), Special Funds from the Chongqing Postdoctoral Science Foundation (Xm2015037, Xm2016044), Key research for Humanities and social sciences of Ministry of Education (14JJD880009). http://fcon_1000.projects.nitrc.org/indi/retro/sald.html

University of Wisconsin, Madison (Birn, Prabhakaran, Meyerand) CoRR sample (UWM): Zuo, X.N., et al. (2014). An open science resource for establishing reliability and reproducibility in functional connectomics. *Scientific data, 1*, 140049. doi: 10.1038/sdata.2014.49 http://fcon_1000.projects.nitrc.org/indi/CoRR/html/uwm_1.html

Wayne State EF Dataset: This dataset was supported by National Institute on Aging grants R01-AG011230, R37-AG011230, R03-AG024630 to Naftali Raz, Ph.D. http://fcon_1000.projects.nitrc.org/indi/retro/wayne_EF.html

Yale Low-Resolution Controls Dataset: Scheinost D, Tokoglu F, Shen X, Finn ES, Noble S, Papademetris X, Constable RT. Fluctuations in Global Brain Activity Are Associated With Changes in Whole-Brain Connectivity of Functional Networks. IEEE Trans Biomed Eng. 2016 Dec;63(12):2540-2549. Epub 2016 Aug 16. http://fcon_1000.projects.nitrc.org/indi/retro/yale_lowres.html

