## Supplementary material for "NOMIS: Quantifying morphometric deviations from normality over the lifetime of the adult human brain": Quality control examples.pdf (from another version of this preprint: version 2, 2022-02-03)

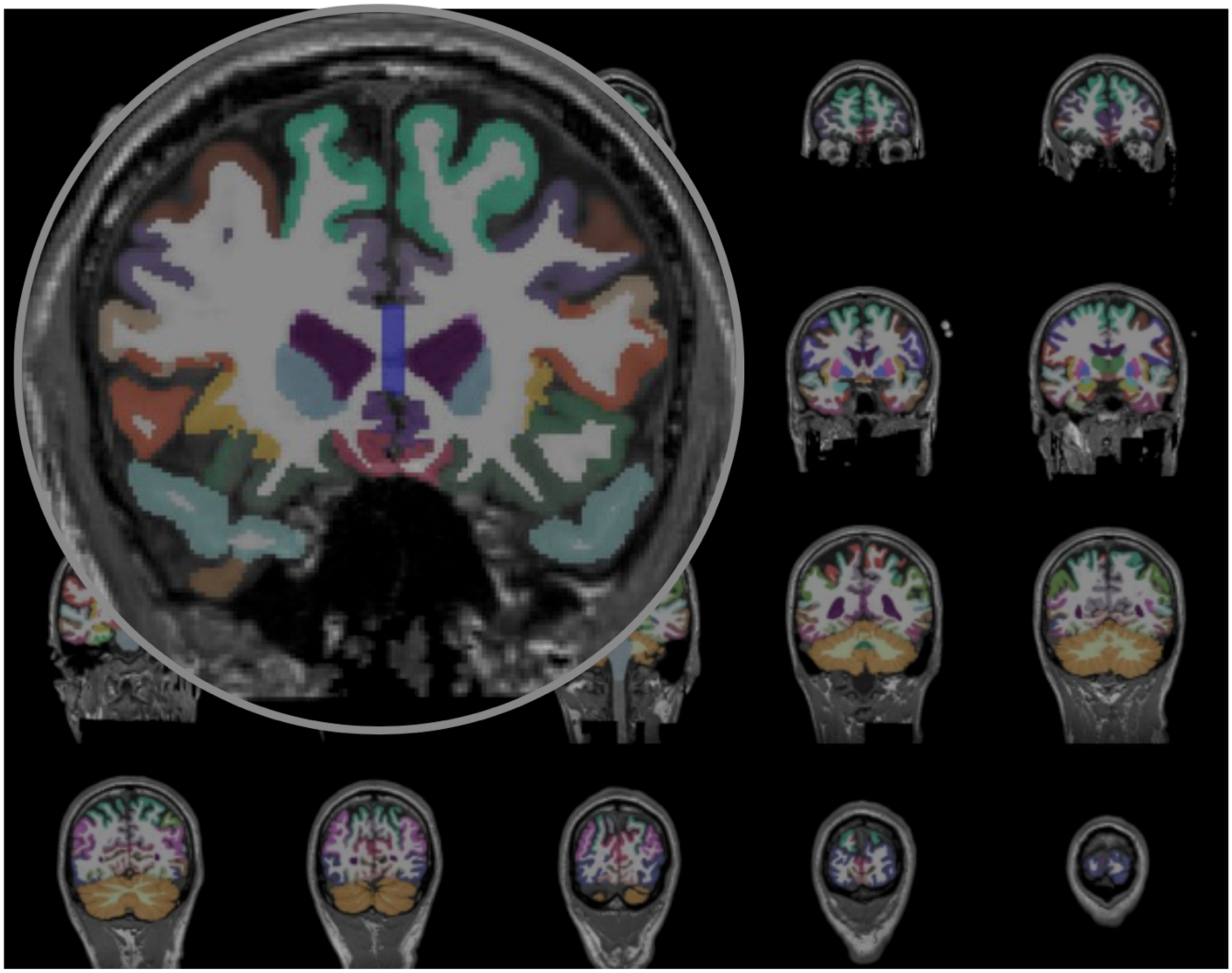

Example of the quality control set up used to verify all images from the normative sample.

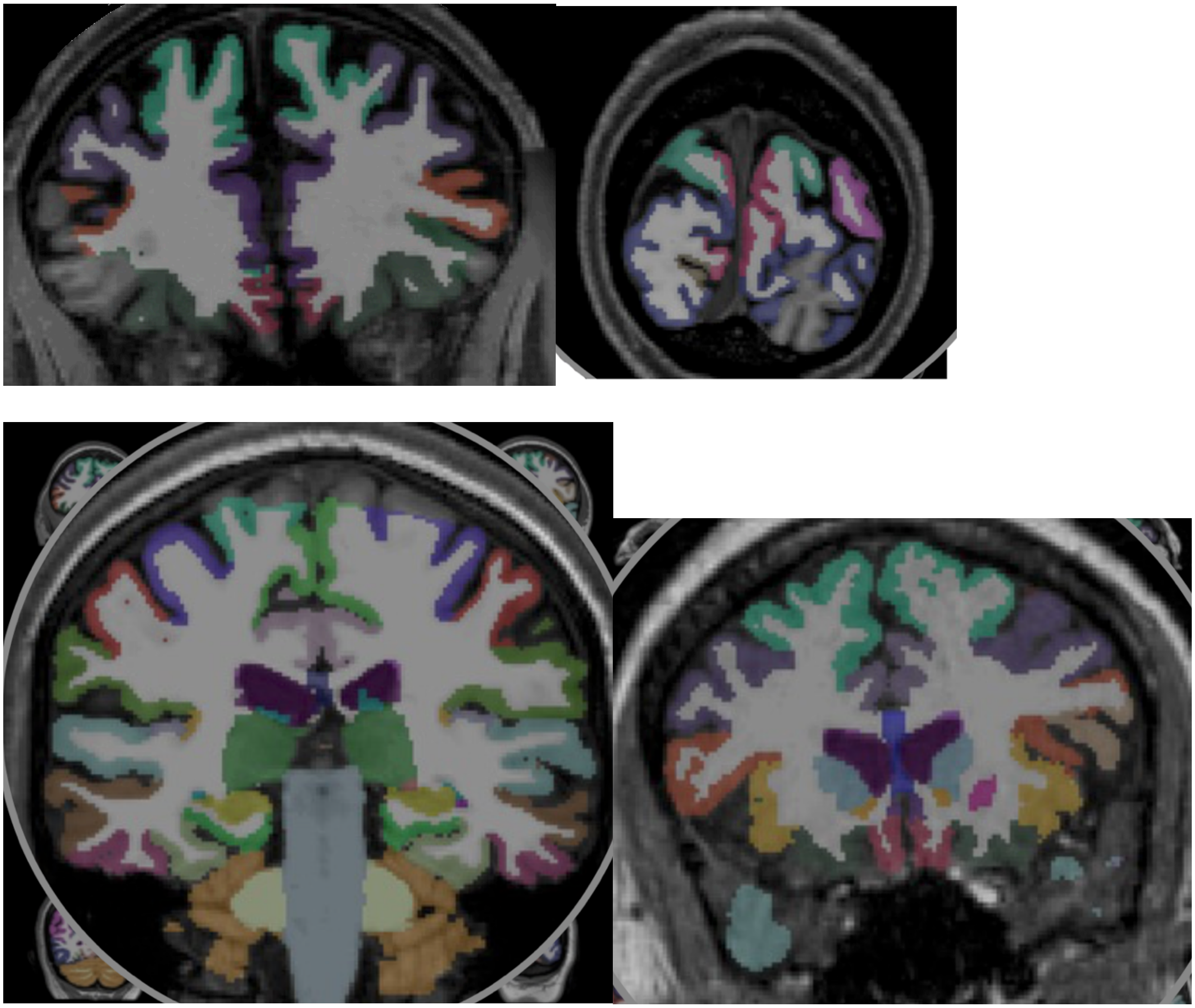

Four examples of eliminated participants because of inadequate segmentation.

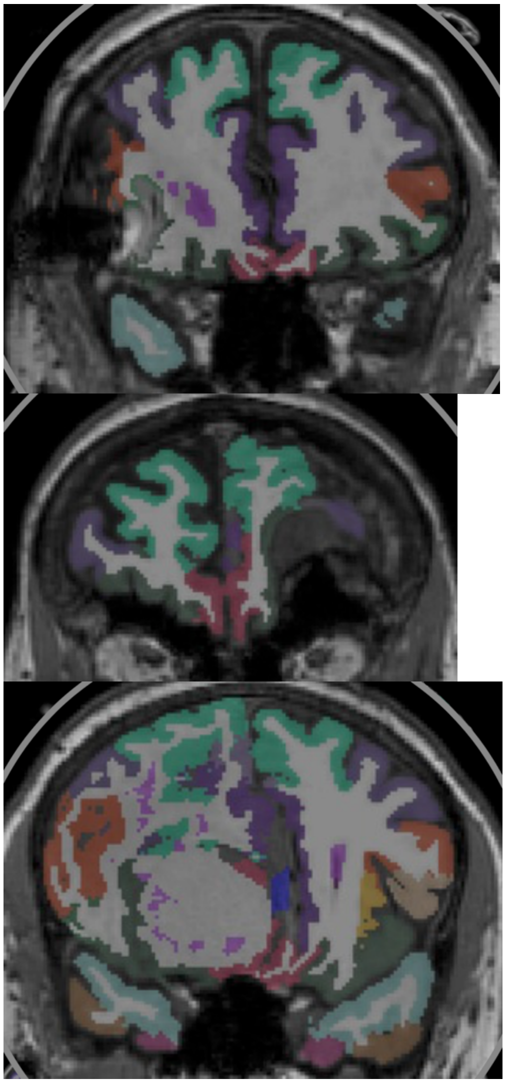

Examples of eliminated participants because of abnormalities.
