## Supplementary figures and images for "NOMIS: Quantifying morphometric deviations from normality over the lifetime of the adult human brain" (from another version of this preprint: version 2, 2022-02-03)

### Supplementary Fig1.tiff

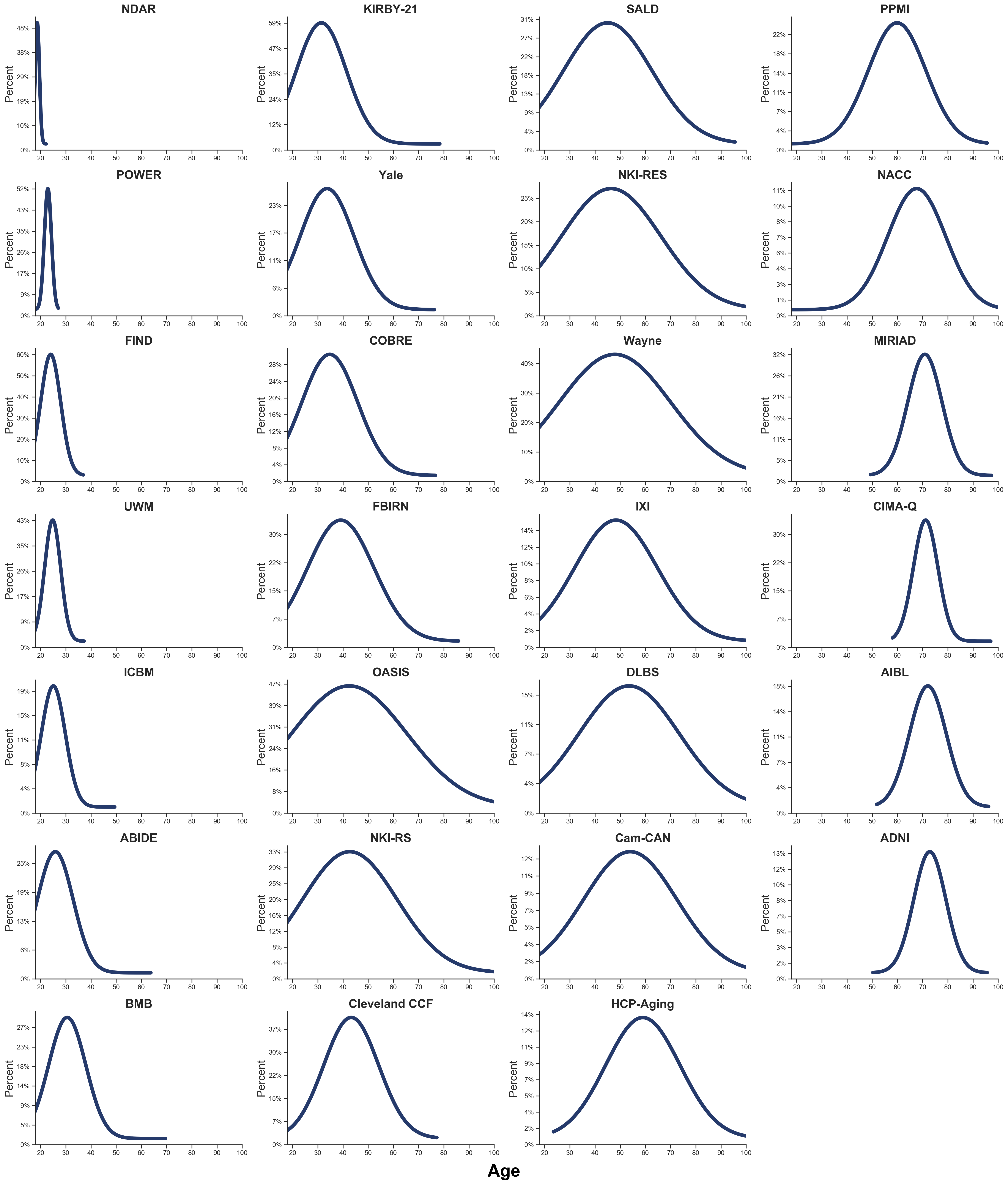

### Supplementary Fig2.tiff

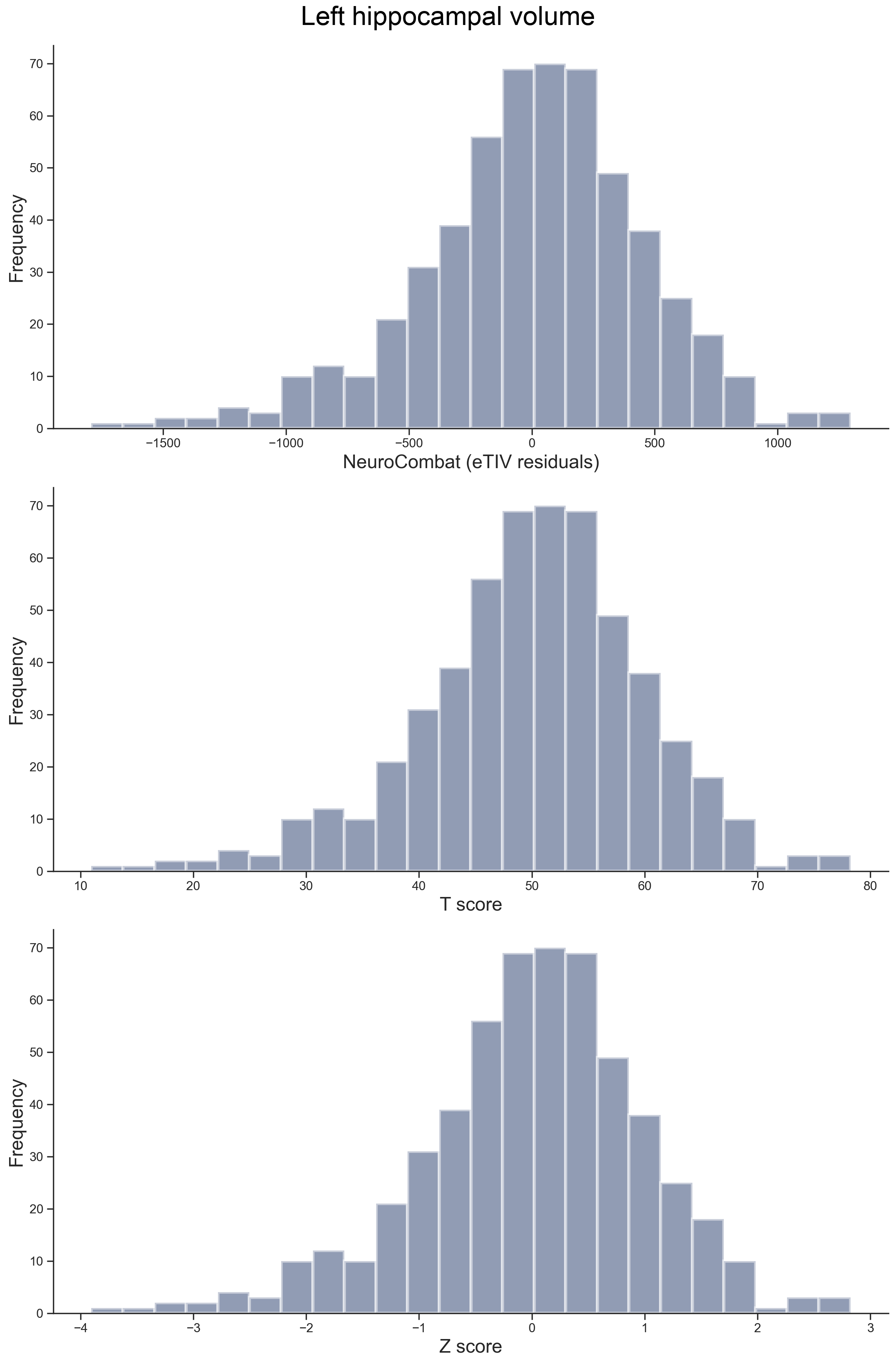

### Supplementary Fig3.tiff

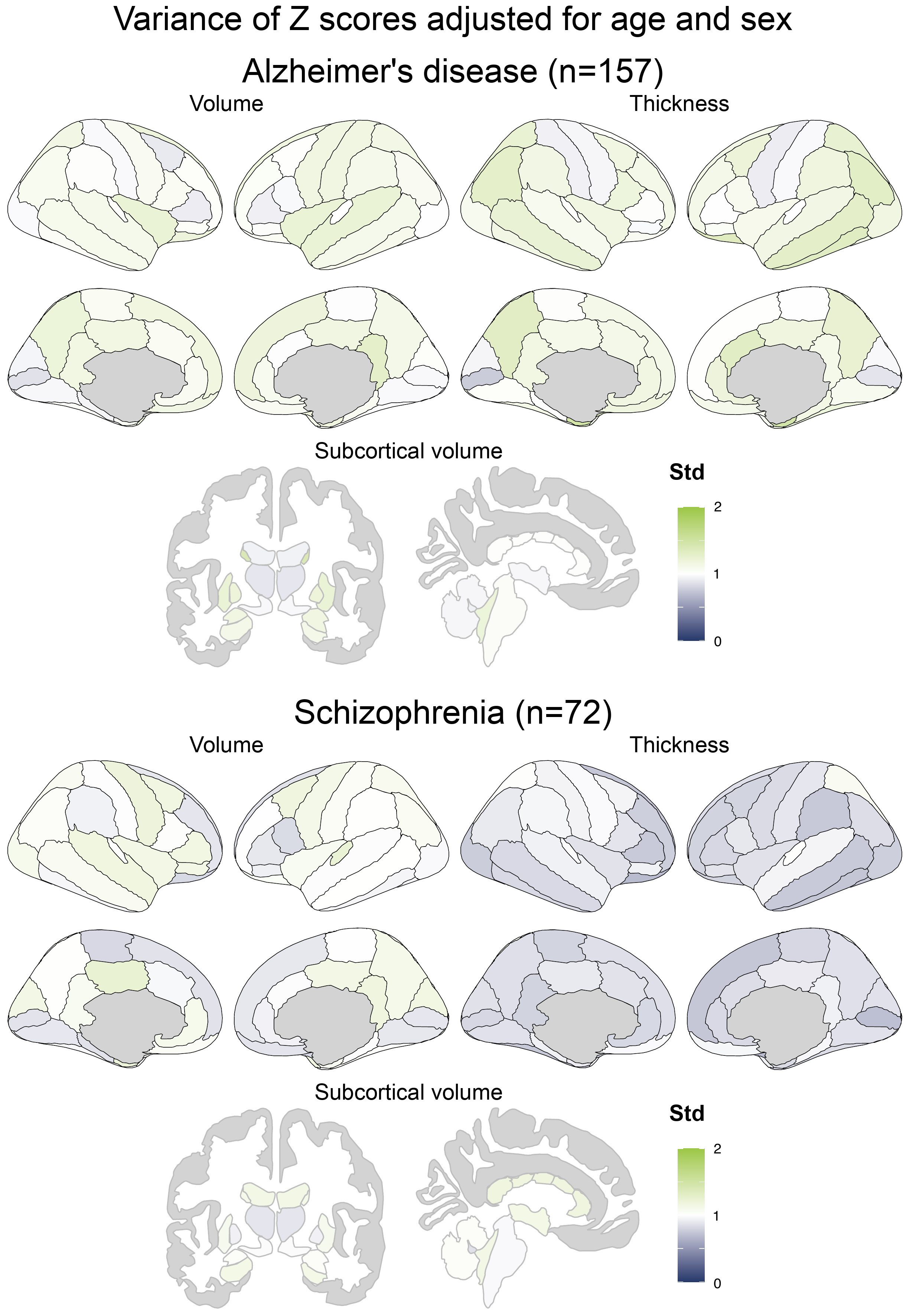

### Supplementary Fig4.tiff

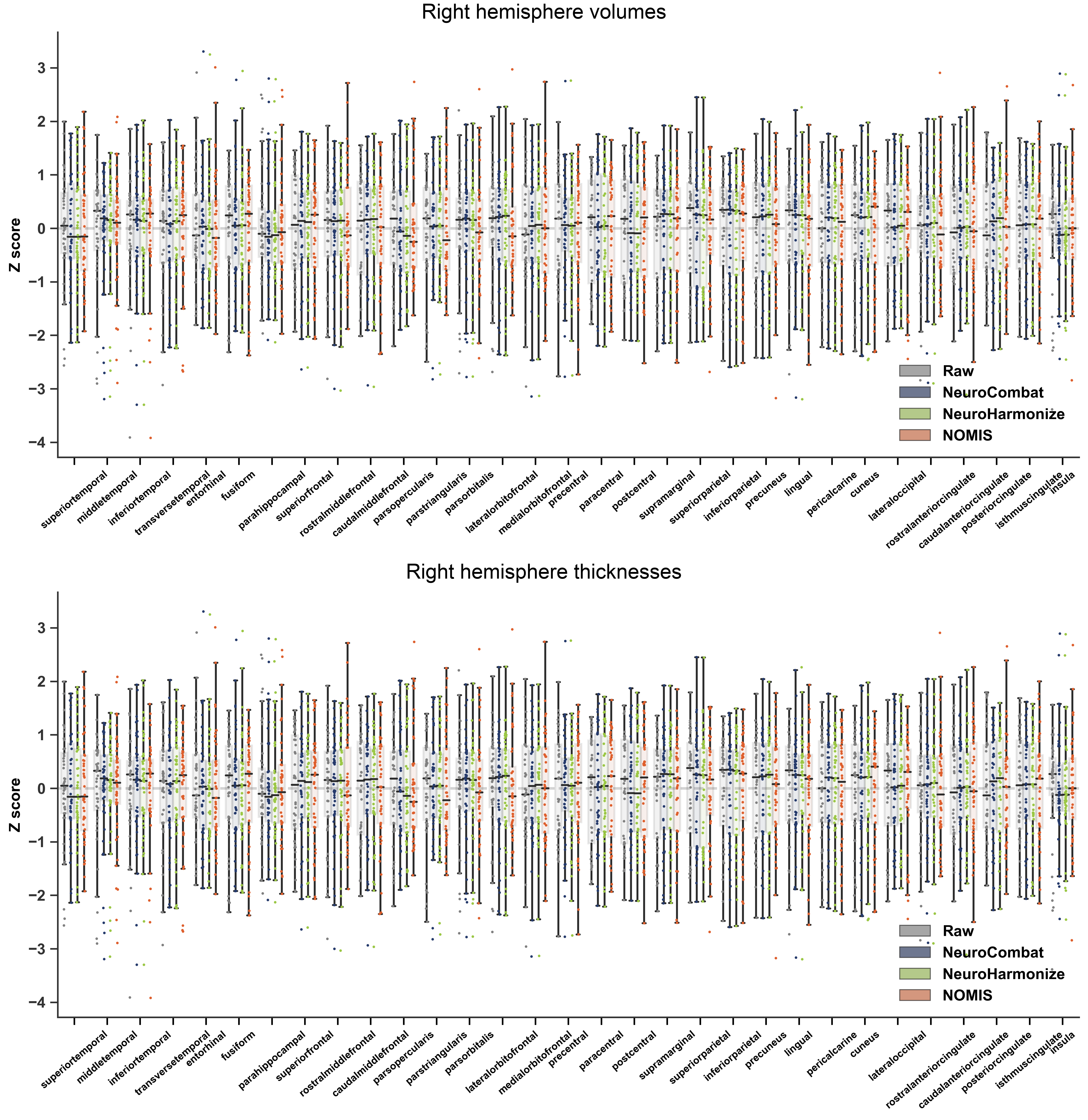

### Supplementary Fig5.tiff

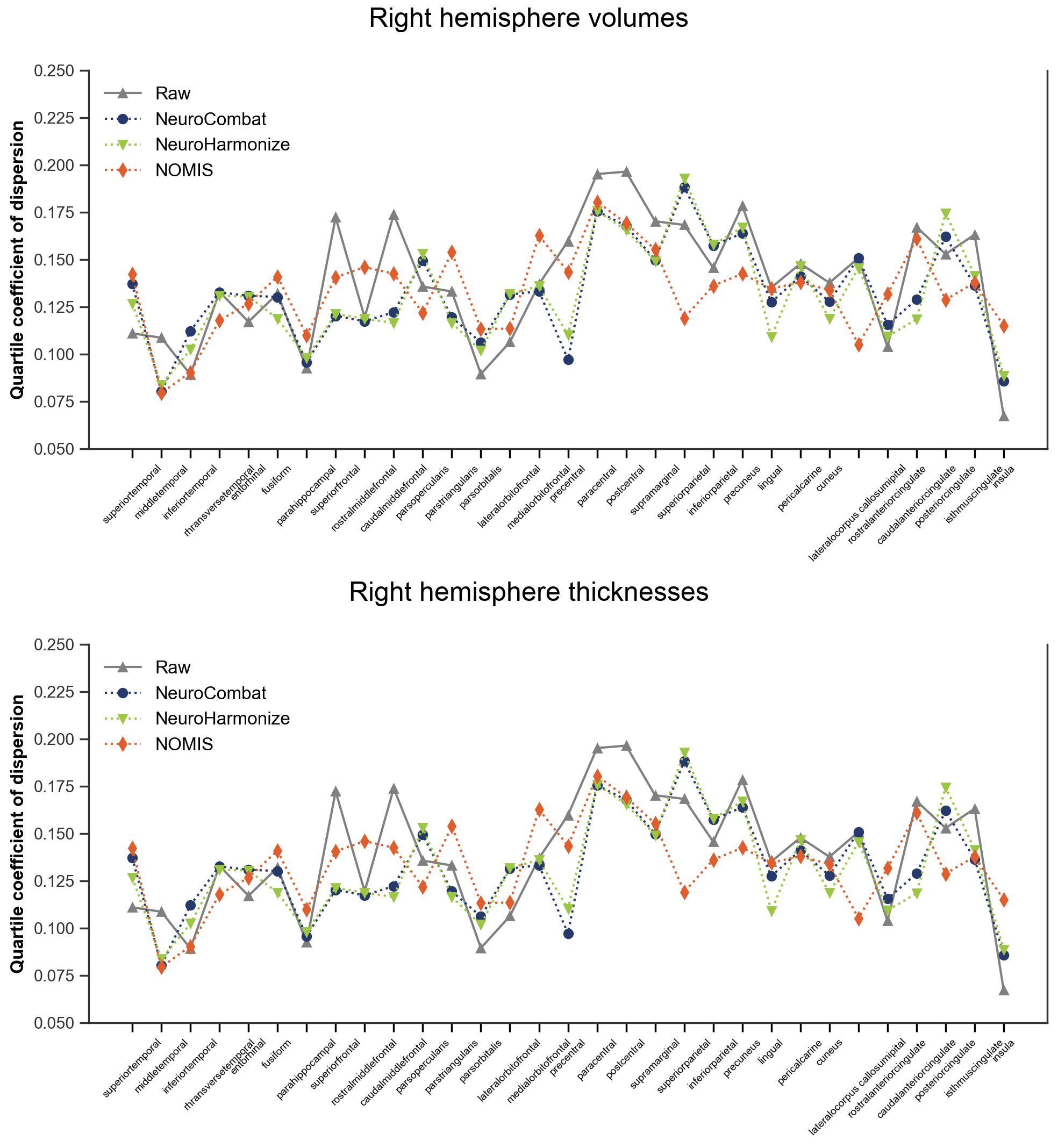

### Supplementary Fig6.tiff

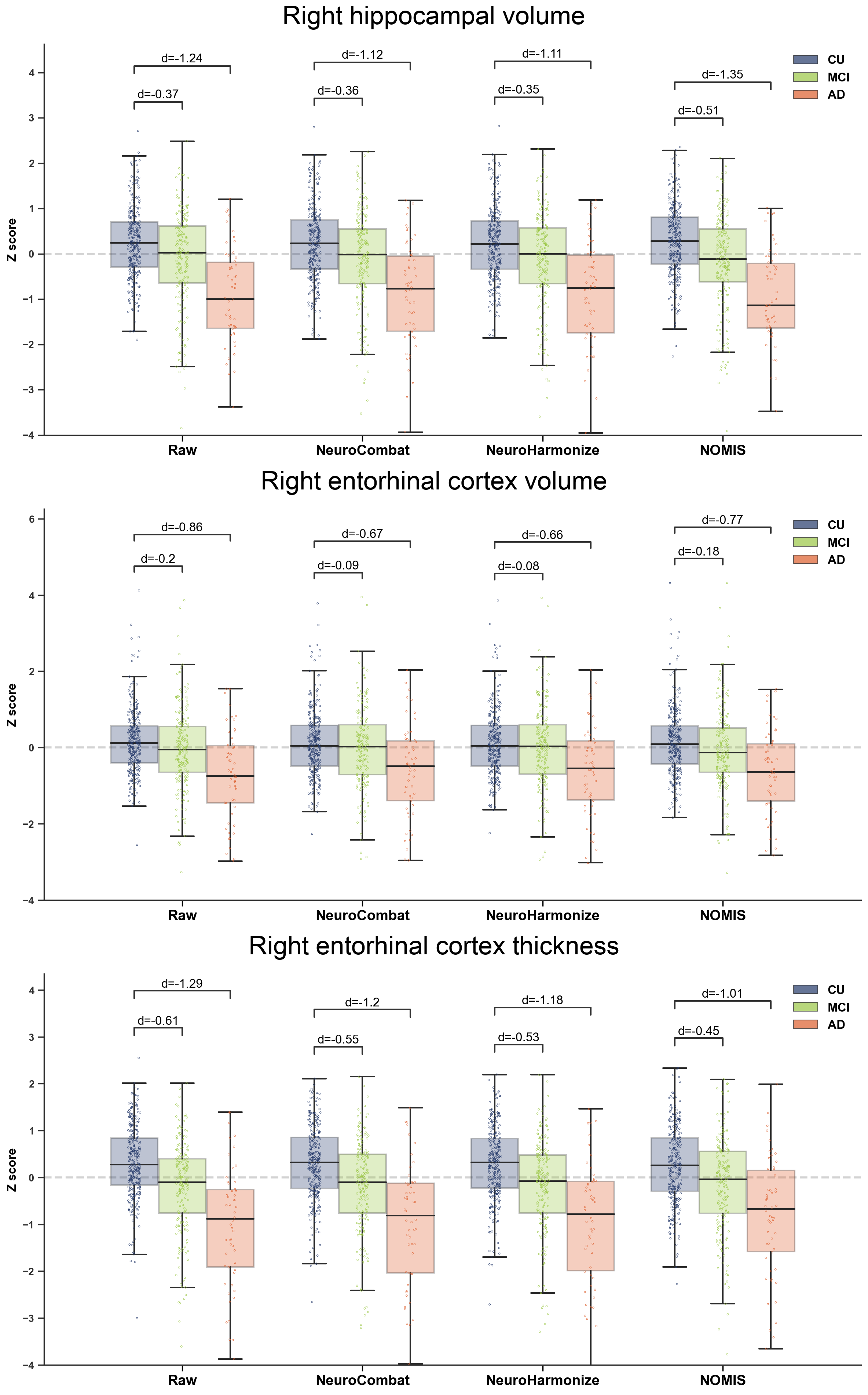
